# A Generalized Higher-order Correlation Analysis Framework for Multi-Omics Network Inference

**DOI:** 10.1101/2024.01.22.576667

**Authors:** Weixuan Liu, Katherine A. Pratte, Peter J. Castaldi, Craig Hersh, Russell P. Bowler, Farnoush Banaei-Kashani, Katerina J. Kechris

## Abstract

Multiple -omics (genomics, proteomics, etc.) profiles are commonly generated to gain insight into a disease or physiological system. Constructing multi-omics networks with respect to the trait(s) of interest provides an opportunity to understand relationships between molecular features but integration is challenging due to multiple data sets with high dimensionality. One approach is to use canonical correlation to integrate one or two omics types and a single trait of interest. However, these types of methods may be limited due to (1) not accounting for higher-order correlations existing among features, (2) computational inefficiency when extending to more than two omics data when using a penalty term-based sparsity method, and (3) lack of flexibility for focusing on specific correlations (e.g., omics-to-phenotype correlation versus omics-to-omics correlations). In this work, we have developed a novel multi-omics network analysis pipeline called Sparse Generalized Tensor Canonical Correlation Analysis Network Inference (SGTCCA-Net) that can effectively overcome these limitations. We also introduce an implementation to improve the summarization of networks for downstream analyses. Simulation and real-data experiments demonstrate the effectiveness of our novel method for inferring omics networks and features of interest.

**Author summary:** Multi-omics network inference is crucial for identifying disease-specific molecular interactions across various molecular profiles, which helps understand the biological processes related to disease etiology. Traditional multi-omics integration methods focus mainly on pairwise interactions by only considering two molecular profiles at a time. This approach overlooks the complex, higher-order correlations often present in multi-omics data, especially when analyzing more than two types of -omics data and phenotypes. Higher-order correlation, by definition, refers to the simultaneous relationships among more than two types of -omics data and phenotype, providing a more complex and complete understanding of the interactions in biological systems. Our research introduces Sparse Generalized Tensor Canonical Correlation Network Analysis (SGTCCA-Net), a novel framework that effectively utilizes both higher-order and lower-order correlations for multi-omics network inference. SGTCCA-Net is adaptable for exploring diverse correlation structures within multi-omics data and is able to construct complex multi-omics networks in a two-dimensional space. This method offers a comprehensive view of molecular feature interactions with respect to complex diseases. Our simulation studies and real data experiments validate SGTCCA-Net as a potent tool for biomarker identification and uncovering biological mechanisms associated with targeted diseases.

## Introduction

### 0.1 Multi-Omics Data Integration

The integration of multiple datasets with the same set of subjects has been actively explored in the field of machine learning, referred to as multi-view machine learning [1]. This approach has a broad range of applications, including clustering subjects based on the consensus of different views [2] and performing image annotation [3]. Furthermore, multi-view machine learning has been used extensively in the biomedical domain [4, 5]. Recent advances in biomedical technologies have enabled the generation of high-throughput data at the molecular level, such as the genome, transcriptome, proteome, and metabolome. Collectively, these datasets are referred to as multi-omics data [6]. Traditionally, each omics dataset has been analyzed separately [7, 8], which can result in the loss of important information about the connections between omics datasets. Multi-omics integration is an alternative approach that identifies shared and complimentary information across different molecular profiles [9]. Integrating multi-omics data can help uncover biological mechanisms and interactions at the molecular level, and has been used for various purposes such as disease subtyping [10, 11], variable selection [12, 13], network analysis [14, 15] and prediction of biomarkers [6, 16, 17].

### 0.2 Multi-View Dimension Reduction

Dimension reduction is one of the more common goals of high dimensional data analysis. For example, non-negative matrix factorization is implemented to cluster multiple data sets [18], and multi-view co-reduction is developed to preserve the locality within the view and the consistency between the views in the lower dimensional embedding of each view [19]. Other dimension reduction methods for multiple data sets are based on Canonical Correlation Analysis (CCA) [20], which seeks to find the linear combination (canonical weight) that maximizes the correlation between two sets of data. Usually, there is more than one solution, which may be referred to as a canonical weight matrix, and thus it is commonly used for dimension reduction by projecting the original data into a shared lower dimensional space between two views (or omics data) with canonical weight matrix. Given two data sets *X*_1_ and *X*_2_, the optimization function is:

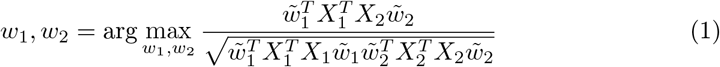

However, this formulation only considers two sets of data and is not generalized to multiple data. A multiple canonical correlation method was developed to maximize the sum of pairwise canonical correlations [21]. Suppose there are *j* = 1, …, *K* views, then the problem can be formulated as follows:

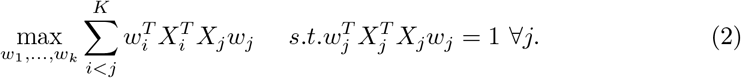

A penalty term can be added to the formulation above to achieve sparsity, which is called Sparse Multiple Canonical Correlation Analysis (SmCCA).

Network analysis for molecular profiles is often used with dimension reduction methods like SmCCA to identify and visualize connections between features [14, 15] for the purpose of inferring underlying biological interactions. In addition to canonical correlation analysis, regression is another approach that is implemented for multi-omics network inference, particularly for gene regulatory networks [22, 23]. However, most methods do not incorporate an outcome or phenotype in the form of a quantitative trait. Canonical correlation analysis-based methods can accomplish this goal by expanding CCA to incorporate a phenotype (*Y*). SmCCNet was proposed to partition omics features into different network modules while considering the phenotype(s) of interest [24] and using a scaled version of SmCCA with the following optimization:

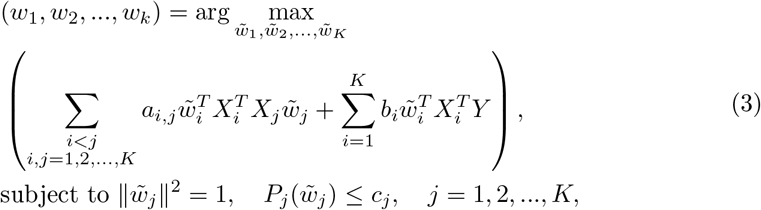

where *a*_*i,j*_ and *b*_*i*_ are scaling factors that place greater importance on pairwise combinations of views. For example, one may want to increase *b*_*i*_ to increase the influence on the phenotype to determine the canonical weights, and *P* (*·*) is the lasso penalty parameter for sparsity [25], but other types of penalties can also be used. The optimal penalty parameter is selected through k-fold cross-validation. After that, the canonical weights can be extracted based on the objective function above to construct an adjacency matrix. Finally, to construct a network, hierarchical clustering can be implemented afterward to extract multiple multi-omics network modules. This method has been applied in various contexts including identifying phenotype-specific miRNA-mRNA and proteomics-metabolomics correlations and networks [26]. However, the existing SmCCNet method can only be adapted to two molecular profiles plus one single phenotype. When extending it to 3 or more omics data types, it can become computationally expensive due to the cross-validation step to find the optimal penalty parameter for each omics data set.

Another multi-omics analysis and network inference method Data Integration Analysis for Biomarker discovery using Latent variable approaches for Omics studies (DIABLO) [27] has a similar formulation to the sparse multiple canonical correlation analysis problems (Eq 3). Although the formulations of SmCCNet and DIABLO are similar, they have some differences: (1) In DIABLO, there is no lasso penalty, but the user can choose how many features to include as nonsparse for each data view instead; (2) DIABLO focuses more on the prediction of phenotype and biomarker discovery, while SmCCNet focuses more on graph learning of the interaction between biomarkers with respect to phenotype; (3) the adjacency matrix of SmCCNet is aggregated through canonical weights obtained from subsampling the data multiple times for a more robust solution, while the adjacency matrix of DIABLO is the subset of correlation matrix with only molecular features selected by the canonical correlation analysis.

### 0.3 Tensor Canonical Correlation Analysis

As reviewed, the dimension reduction methods based on canonical correlation analysis only consider the summation of all pairwise correlation, including methods like SmCCNet and DIABLO. However, for multi-omics data with more than two molecular profiles, the correlation structure can be higher-order rather than pairwise, where higher-order correlation is defined as the simultaneous correlation among 3 or more features. This type of higher-order correlation can be captured by pairwise correlation if and only if all pairwise correlations are strong. However, in multi-omics data, it may not be the case that higher-order correlations also have strong pairwise correlations.

A particular variant of the multiple canonical correlation analysis extends the pairwise relationship to a higher-order relationship by maximizing the tensor canonical correlation [28], which captures higher-order correlations among multiple data sets and projects them into a shared lower dimensional embedding. A further extension to this tensor canonical correlation analysis (TCCA) is to combine deep learning with tensor canonical correlation analysis to maximize the higher-order correlation between views in the non-linear lower-dimensional space for dimensional reduction [29] to allow for non-linearity.

### 0.4 Contributions

As reviewed, SmCCA captures pairwise correlation, while TCCA captures higher-order correlation. However, TCCA only captures the highest order correlation, but not all types of lower-order correlation structures of interest. In addition, sparsity is not considered in TCCA, which prevents the model from focusing only on the most relevant information in the data. Furthermore, TCCA has a high memory and computation cost, which is inefficient for high-dimensional omics data. Therefore, in this work, we present a new method for identifying multi-omics phenotype specifics network that is based on TCCA but with significant extensions by including correlations of higher and lower order simultaneously (Figure 1) and by incorporating sparsity. This new method is called Sparse Generalized Tensor Canonical Correlation Analysis for Network Analysis (SGTCCA-Net). In particular, we avoid repeatedly applying TCCA to each correlation structure, but instead use a simultaneous algorithm that accounts for all the correlation components.

**Fig 1.**
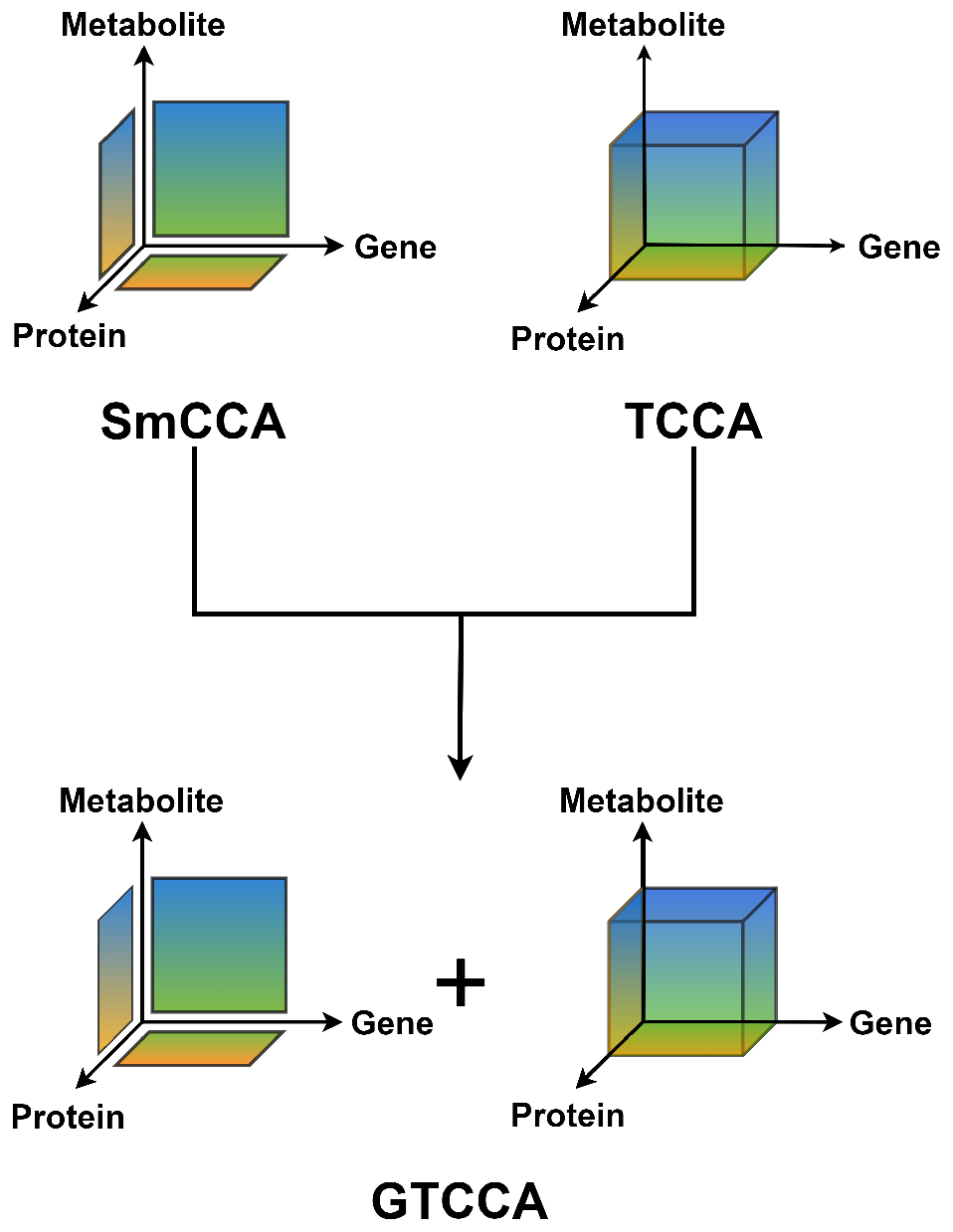
Comparison between SmCCA, TCCA, and GTCCA. Visualization of the comparison between Sparse multiple Canonical Correlation Analysis (SmCCA), Tensor Canonical Correlation Analysis (TCCA), and Generalized Tensor Canonical Correlation Analysis (GTCCA). SmCCA only captures pairwise correlations (e.g., genes and proteins) and TCCA only captures the highest order correlations (e.g., gene, protein and metabolites),. GTCCA considers the combination of highest order correlations ((TCCA) and lower-order correlations (SmCCA).

In this novel pipeline, we introduce multiple major contributions: New definition of higher-order correlation compared to the original TCCA, which better measures higher-order correlation without interpreting the directionality and avoids effect cancellation issues; The development of Sparse Generalized Tensor Canonical Correlation Analysis (SGTCCA) with consideration of higher-order correlation, flexible design of correlation structures, and biased subsampling algorithm that ensures sparsity and computational efficiency; Implementation of network pruning and summarization algorithms to keep only the most important molecular features. Our pipeline first performs feature selection with SGTCCA, which extracts molecular features that are involved in one or more specified higher/lower-order correlation structures. Then an adjacency matrix between selected features is constructed to identify the inter-molecule relationships and collapse all higher/lower-order correlations into the two-dimensional space. Afterward, a network pruning algorithm is implemented to the adjacency matrix with the PageRank algorithm [30] NetSHy summarization score [31] to reduce the size of the network and include only the most relevant molecular features, yielding the final multi-omics network. Details of the pipeline are described below

## Materials and methods

### Summary of Pipeline Workflow

The end-to-end pipeline for SGTCCA-Net (Figure 2 and Algorithm 1) inputs the multi-omics data 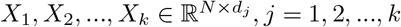, each with dimension *d*_*j*_, and outputs subnetwork adjacency matrix 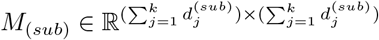, where 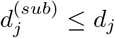 for all *j* = 1, 2, …, *k* is the number of features belong to the *j*th molecular profile. There are two major contributions in this paper: (1) the development of Sparse Generalized Tensor Canonical Correlation Analysis that combines higher-order and lower-order correlation of interests and guarantees high computational efficiency, and (2) a novel network pruning algorithm based on the network summarization score.

**Fig 2.**
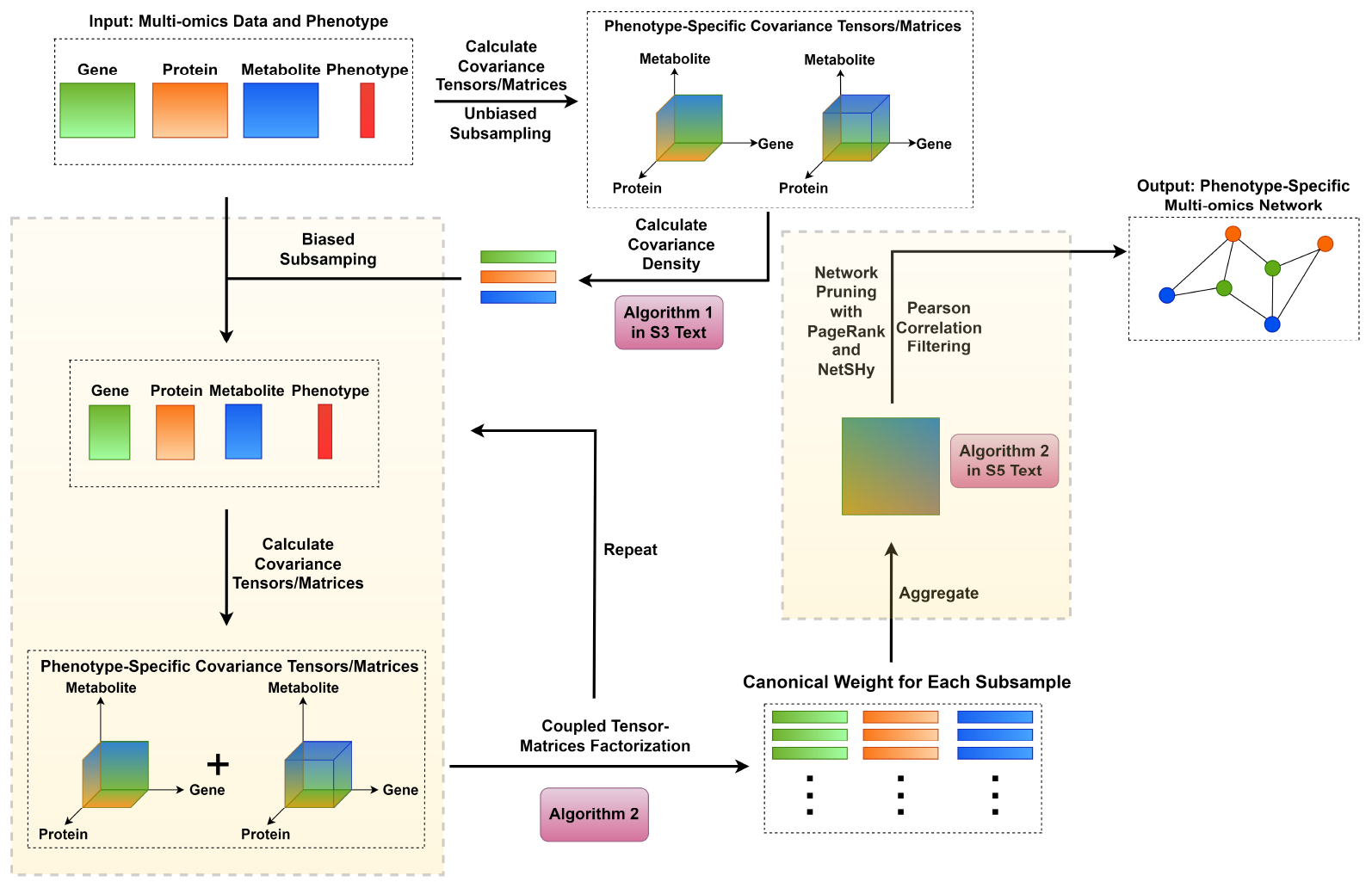
SGTCCA-Net workflow. Workflow of SGTCCA-Net pipeline for multi-omics network inference. It consists of three core steps: higher-order correlation extraction, affinity matrix construction, and network trimming. Biased subsampling involves calculating the covariance tensor/matrix density with respect to certain molecular profiles, which is also the importance/probability vector when randomly sampling features from each molecular profile.

#### Algorithm 1

SGTCCA-Net End-to-End Pipeline Algorithm.

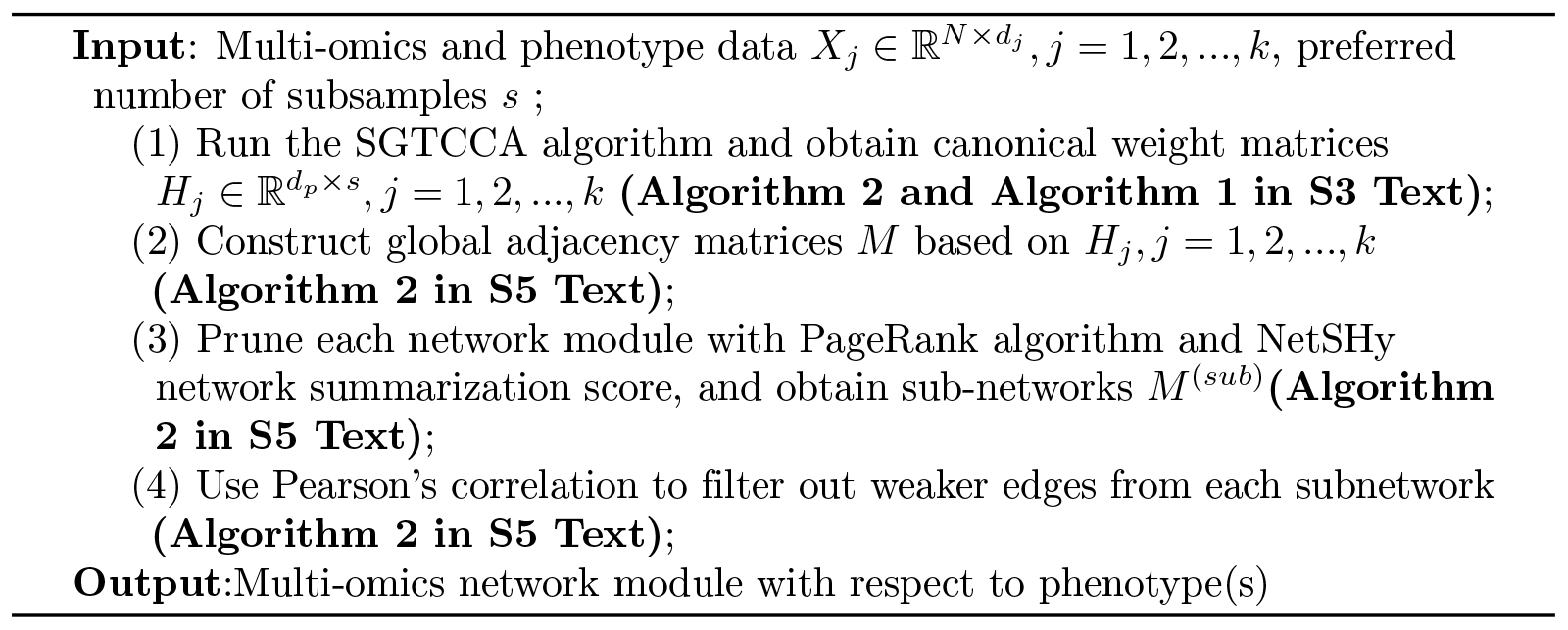

### 0.6 Generalized Tensor Canonical Correlation Analysis

#### 0.6.1 New Tensor Canonical Correlation Analysis

The original TCCA algorithm has two major issues: (1) the directionality (positive/negative) of higher-order correlation is not interpretable, especially when there are more than three datasets, (2) the correlation effect will be cancelled for odd number of datasets (see S1 Text for detail). Therefore, we addressed these limitations by proposing a new tensor canonical correlation analysis formulation. Let *z*_1_, *z*_2_, …, *z*_*k*_ *∈ ℝ*^*n×*1^ be *k n ×* 1 dimensional vectors that are centered and scaled, and *∈* ℝ^*n×*1^ is the all-one *n ×* 1 vector, then the higher-order correlation between these **0** vectors can be defined as:

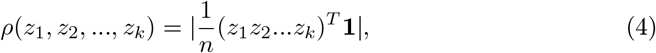

where *z*_1_*z*_2_…*z*_*k*_ is the element-wise multiplication between *k* vectors (assuming *k* is even). The higher-order correlation calculation method is slightly different for the odd number of views to avoid effect cancellation (see S1 Text for detail). Eq 4 calculates the higher-order correlation between vectors, but if the higher-order correlation between matrices (e.g. each matrix is an omics dataset) needs to be calculated, we can avoid processing multiple loops to calculate the higher-order correlation for all combinations of features through a different equation. Suppose that there are *k* data in total, denoted by 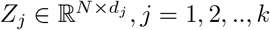, which are centered and scaled. Let 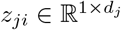 be the vector for subject *i* in the *j*th view. For each view, there are *N* observations and *d*_*j*_ features, then the covariance tensor 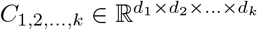 for *k* views can be denoted by:

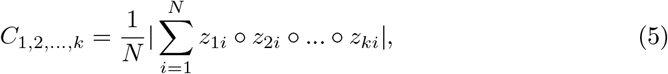

where *∘* denotes the outer product. It can be shown that each entry of *C*_1,2,…,*k*_, denoted by *C*_1,2,…,*k*_(*j*_1_, *j*_2_, …, *j*_*k*_), can also be calculated through Eq 4 (see S1 Text for more details). If the canonical weights for each view are 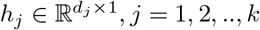, then our new tensor canonical correlation analysis is formulated as follows:

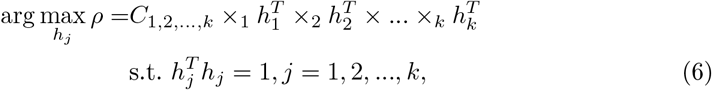

where the *i*-mode product of *C*_1,2,…,*k*_ and 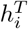, denoted by *C*_1,2,…,*k*_ *×*_*i*_ *h*_*i*_ is defined as follow:

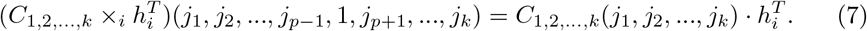

It has been shown that the optimization problem above is equivalent to the following form [32]:

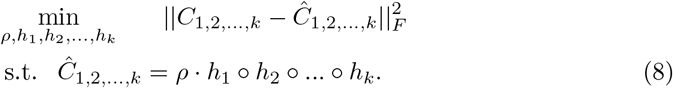

In this optimization form, the canonical weights *h*_*j*_ are solved using either a gradient-based method or alternating least squares [33].

#### 0.6.2 Generalized Tensor Canonical Correlation Analysis Combining Higher and Lower-Order Correlations

##### Algorithm 2

Generalized Tensor Canonical Correlation Analysis Algorithm.

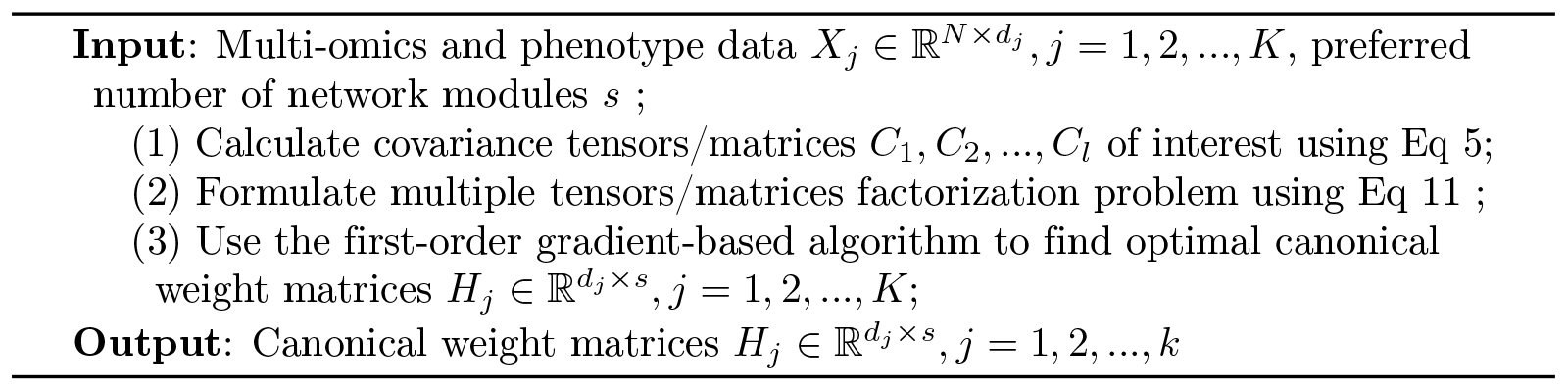

The section above illustrates TCCA assuming that only a single full covariance tensor is used and is not capable of adding more covariance structures. For example, with the presence of transcriptomic, proteomics, metabolomics, and phenotype data, we may also be interested in the 3-way or pairwise correlation with respect to phenotype in addition to the full 4-way correlation structure. Therefore, to incorporate multiple correlation structures of interest, we extended our new TCCA method and developed the Generalized Tensor Canonical Correlation Analysis (GTCCA).

Let *S*_*m*_ = *{*(*m*_1_, …, *m*_*m*_) : *m*_*i*_ *∈ {*1, …, *k}, m*_1_ ≠*m*_2_ *≠* … *≠ m*_*m*_*}* be all possible combinations of *k* choose *m*, let *S*_*m*_(*i*) be the *i*th element in the set. For instance, with *k* = 3 and *m* = 2, it would be the set of *{S*_2_(1) = (1, 2), *S*_2_(2) = (1, 3), *S*_2_(3) = (2, 3)*}*, then the GTCCA can be defined as:

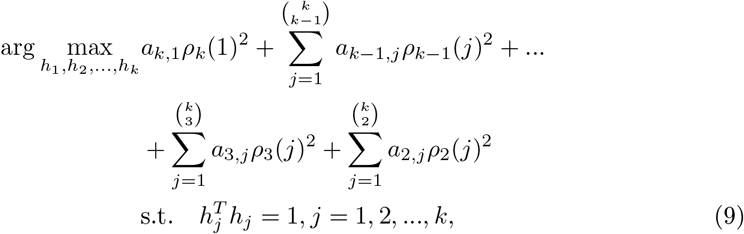

where 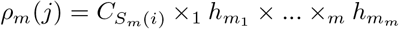, where (*m*_1_, *m*_2_, …, *m*_*m*_) = *S*_*m*_(*i*). Compared to TCCA, this design allows flexibility with respect to a specific experimental design of interest by allowing a portion of *a*_*i,j*_ to be 0. For example, if the investigator is not interested in a 3-way higher-order correlation between omics data 1,2, and 3, then the scaling factor associated with *ρ*_1,2,3_ can be set to 0. In addition, the scaling factor *a*_*i,j*_ can be set to values other than 1 to prioritize certain correlation structures. Eq 9 cannot be directly optimized with the existing gradient-based method, and thus we found the problem equivalency as follows (proof in S2 Text):

##### Theorem 1

*Let* 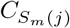 *be the covariance tensor of view with* (*m*_1_, …, *m*_*m*_) = *S*_*m*_(*j*) *such that* 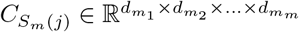, *If the optimization goal is formulated as follows:*

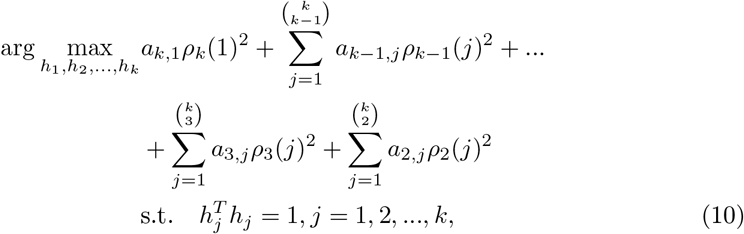

*where* 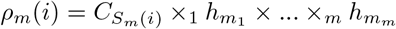 *for all m* = 1, 2, …, *k, then the optimization problem above is equivalent to the following:*

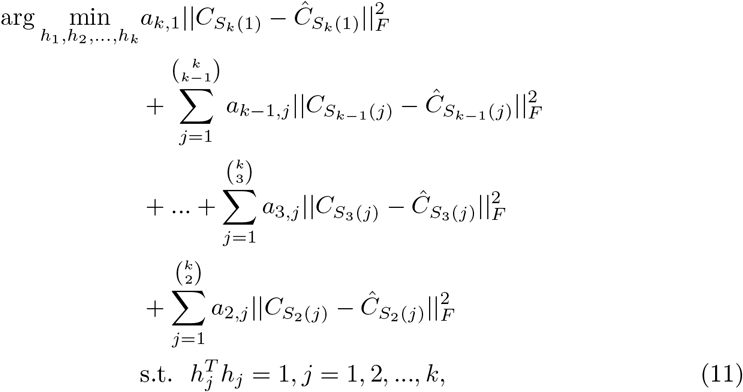

*where* 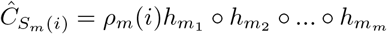 *is the rank-1 approximation of* 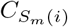.

Since the original problem is shown to be equivalent to the multiple tensor factorization problem, it can be solved iteratively using any first-order gradient-based algorithm. We choose the non-linear conjugate gradient algorithm with the Dai/Yuan update and restart [34, 35].

### 0.7 Multi-Omics Data Example

In this section, we demonstrate how the first-order gradient of Eq 11 can be calculated with a multi-omics data example. Suppose there are three types of molecular profiles: transcriptomics (tr), proteomics (pr), and metabolomics (me), and phenotype (ph) data.

Define these data as *X*_*tr*_, *X*_*pr*_, *X*_*me*_ and *Y*_*ph*_. Using GTCCA to find the phenotype-related correlation structure, the optimization problem is given by:

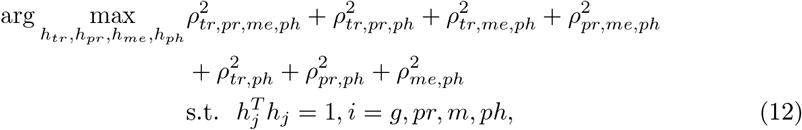

where *ρ*_*tr,pr,me,ph*_ = *C*_*tr,pr,me,ph*_ *×*_1_ *h*_*tr*_ *×*_2_ *h*_*pr*_ *×*_3_ *h*_*me*_ *×*_4_ *h*_*ph*_, and the other correlation components are also in the same form. By Theorem (1), optimizing this objective function is equivalent to:

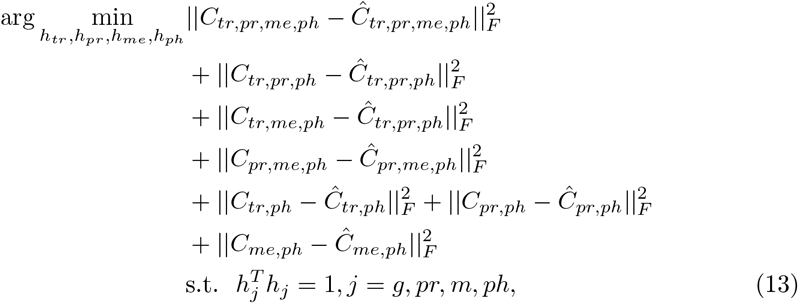

where 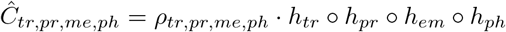 represents the rank-1 approximation of the covariance tensor, and other components are similar. Below is the example gradient calculation for the transcriptome *h*_*tr*_,

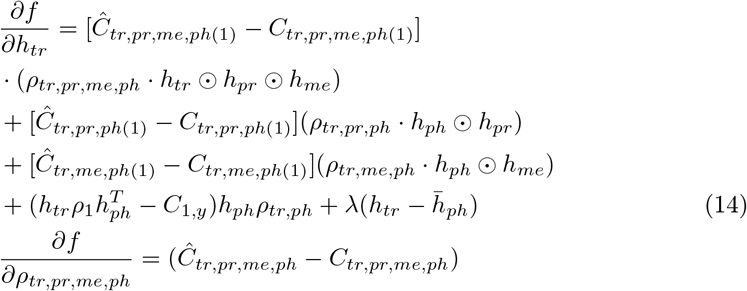

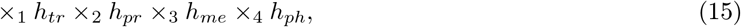

where *⊙* is the Khatri-Rao product, given two matrices 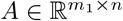 and 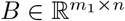, and let *⊗* denote the Kronecker product between two vectors, the Khatri-Rao product is given by:

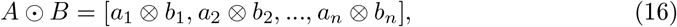

*C*_*tr,pr,me,ph*(*i*)_ is the mode-*i* matricization of tensor *C*_*tr,pr,me,ph*_, and “mode” stands for the index of dimension in tensor data, and “mode-*i*” means the *i*th dimension of the tensor data. This is a way to matricize the tensor by mapping the elements of the tensor into a matrix by arranging the mode-i fibers (think of it as the higher-order rows and columns) so that they become the columns of [*C*_*tr,pr,me,ph*_]_(*i*)_. The gradient for other *h*s and *ρ*s can be calculated in a similar way. After taking the gradient, the next step is to concatenate all of the gradients into a long vector. All first-order gradient-based methods can be used for optimization, and the nonlinear conjugate gradient method is chosen to solve for the optimal canonical weights [36].

### 0.8 Sparse Generalized Tensor Canonical Correlation Analysis

Common methods for ensuring sparsity in statistical models include using penalty-based techniques such as lasso [25] and elastic net [37]. However, applying these methods to GTCCA in practice can be computationally intensive, as they require tensor computation on the original covariance tensors, which can potentially cause memory issues. To address this problem, adapting the idea from Turbo-SMT [38], we use the biased subsampling method to guarantee sparsity (the details of the algorithm setup are shown in Algorithm 1 in S3 Text.). This method is both accurate and efficient, reducing the tensor factorization problem by 1000 times. The core concept of this method is to subsample features based on the covariance density value calculated for each feature. If a feature is more likely to be connected to other features or phenotypes, then it is more likely to be selected in the biased subsampling phase. Details of the calculation of the covariance density are given in S4 Text.

In practice, multi-omics data are often high-dimensional, which poses challenges in terms of memory and computation when computing covariance tensors. Our preliminary findings indicate that in GTCCA optimization, even a covariance tensor of size 1000 *×* 1000 *×* 1000 requires more than 16 GB of RAM, leading to memory explosion. Although SGTCCA reduces the GTCCA memory requirement by feature subsampling, it still requires the calculation of covariance density vectors, which also involves the computation of the full covariance tensor. Therefore, to further improve the algorithm efficiency, when calculating the covariance density vectors, we used the unbiased feature subsampling approach to approximate each covariance tensor. In each subsampling iteration of our method, a small, randomly selected subset of features from each molecular profile is used. For these subsets, we calculate the covariance density based on the provided correlation structures. This process is repeated across multiple iterations. The final estimated covariance density vector is obtained by averaging the covariance densities from all subsamples. As a general rule for setting parameters, a lower subsampling percentage (e.g., 10%) requires a greater number of iterations to achieve reliable results, while a higher subsampling percentage (e.g., 70%) typically requires fewer iterations.

To ensure the consistency of canonical weight vectors from each subsample, a portion of features from each dataset are preselected and shared across different subsamples. For instance, for each subsample, we can have 8% of the features shared across all subsamples and 2% of the distinct features. A general recommendation is to set the total percentage of featured subsampled to be between 10% and 20% to ensure both computational efficiency.

### 0.9 Network Construction and Pruning

After obtaining the canonical weight through the SGTCCA, the next step is to construct an adjacency matrix. The adjacency matrix construction process is the same as SmCCNet, and the detailed algorithm is illustrated in Algorithm 2 of S5 Text. The general idea is to take the outer product between the concatenated canonical weight and itself. However, even though the adjacency matrix is sparse, it may still contain features/nodes that are less associated with other features/phenotypes. Therefore, we prune the global network with the PageRank algorithm [39] and the NetSHy network summarization score [31]. The original PageRank algorithm is widely used to rank web pages according to their importance. Its application to networks (adjacency matrix) is to count the number and strength of the edges for each node to determine the importance of each node. The NetSHy summarization score is the principal component score that takes into account the graph/network topology. We chose the NetSHy summarization score rather than the regular principal component analysis to prioritize the contribution of nodes with high network connectivity. The network pruning algorithm ensures both a high correlation of the summarization score with the phenotype and high network connectivity. Step-by-step details of the algorithm can be found in S5 Text.

The algorithm above provides node-wise network pruning, but the pruned sub-networks will be highly densely connected, and it requires edge pruning. Therefore, we calculate the between-node Pearson’s correlation and use it to filter out edges that connect to two weakly or non-correlated nodes, which improves the final network visualization.

### 0.10 Computational Complexity

Suppose there are *k* omics data, each with *d*_1_, *d*_2_, …, *d*_*k*_ features, the common subsampling proportion is *p*_*c*_ and the distinct subsampling proportion is *p*_*d*_ (see S3 Text for details). Define *p* = *p*_*c*_ + *p*_*d*_ *∈* [0, 1], then the new tensor canonical correlation analysis defined in this paper has the space complexity of *O*(*d*_1_*d*_2_…*d*_*k*_), while SGTCCA has the space complexity of *O*(*d*_1_*d*_2_…*d*_*k*_*p*^*k*^). The computational cost per iteration for new tensor canonical correlation analysis is *O*(*d*_1_*d*_2_…*d*_*k*_*s*), while for SGTCCA it is *O*(*d*_1_*d*_2_…*d*_*k*_*p*^*k*^*s*). Since 0 *< p <* 1, SGTCCA has a more efficient space complexity and computational cost than the new TCCA.

## Experiments

### 0.11 Simulation Study

We simulated multi-omics data using independent latent factors to represent various correlation structures [40] (Table 1). We aimed to compare the performance of SGTCCA-Net with other multi-omics network inference pipelines. Since the comparison relies on the adjacency matrix, it is necessary that all methods have an available adjacency matrix. We generated three datasets for molecular profiles, termed omics 1, omics 2, and omics 3, along with a quantitative phenotype. To mimic the true multi-omics correlation structure in most scenarios (the strong connection between omics but the weak connection between omics and phenotype), we imposed weaker noise on the omics data and stronger noise on the simulated phenotype data, and all the random noise are generated from Gaussian distribution. This simulation study is designed to evaluate the accuracy of different methods in identifying omics features associated with both higher-order and lower-order phenotype-specific correlation structures, and to assess the robustness of the model in handling noisy or highly skewed omics data. To this end, we introduced three latent factors simulation strategies: multivariate normal latent factors, highly right-skewed latent factors (Fleishman power transformation [41]), and noisy latent factors (multivariate normal distribution with noise). Under each simulation strategy, we proposed three distinct correlation structure setups: one with all possible phenotype-specific higher-order/lower-order correlations, one with only phenotype-specific four-way correlations, and one with all phenotype-specific correlation structures except for the 4-way correlations. These signal correlation structures are simulated by latent factors (red in Table 1). Furthermore, we also simulated nonphenotype-specific correlation structures to interfere with signal feature identification and test the robustness of each method (black in Table 1).

**Table 1.**
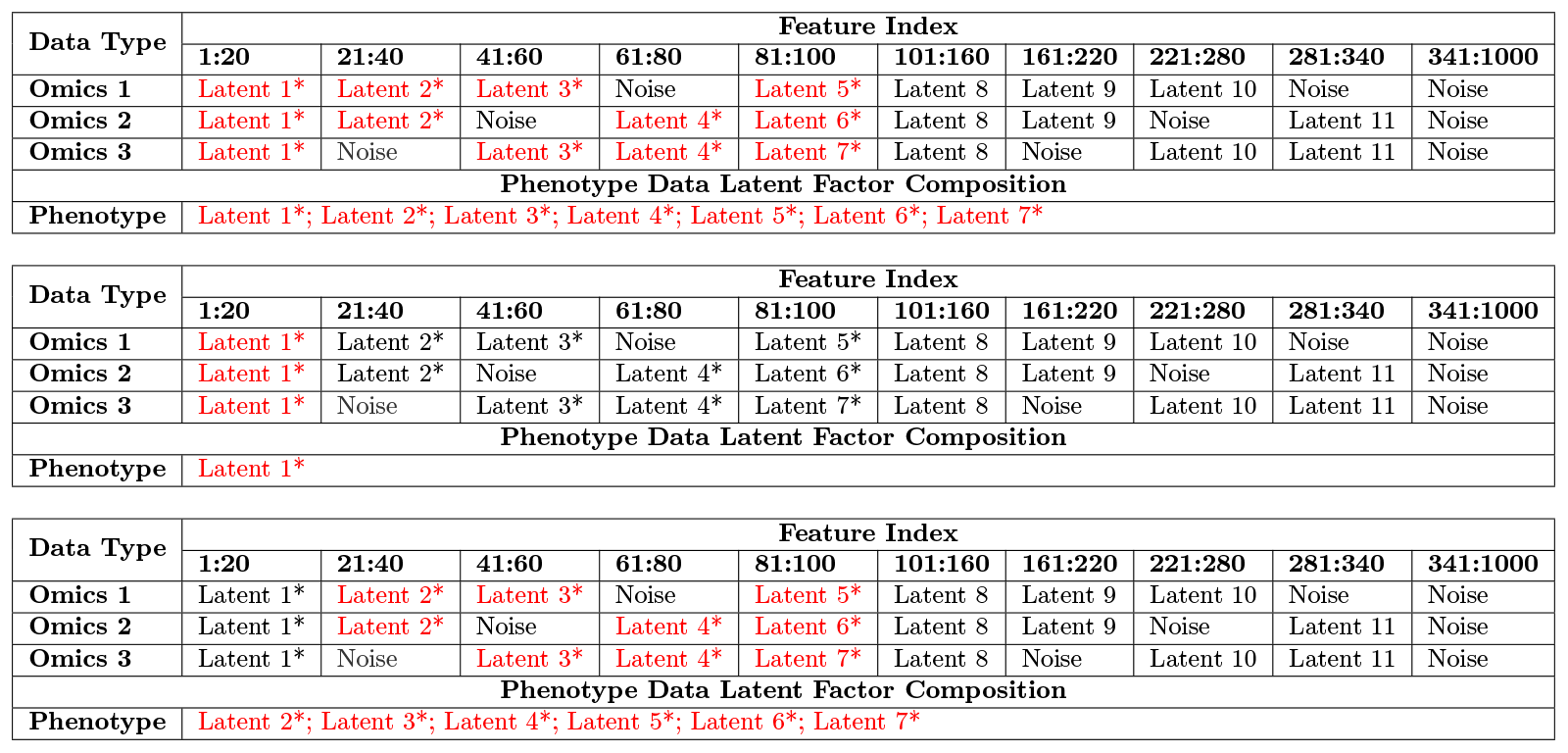
Simulated multi-omics data correlation structure for cases 1-3. Red and “*” mean that features that are simulated with this latent factor are considered signal features. In addition to the existing latent factors and random noise, as shown in the table, additional random noise will be added to all simulated molecular features and phenotype data. The first table is for simulation case 1, where all types of phenotype-specific correlation structures are simulated and considered signal; the second table is for simulation case 2, where only 4-way phenotype-specific correlation structures are simulated and considered signal; the third table is for simulation case 3, where all 3-way, pairwise phenotype-specific correlation structure is simulated and considered signal.

| Data Type | Feature Index |  |  |  |  |  |  |  |  |  |
| --- | --- | --- | --- | --- | --- | --- | --- | --- | --- | --- |
|  | 1:20 | 21:40 | 41:60 | 61:80 | 81:100 | 101:160 | 161:220 | 221:280 | 281:340 | 341:1000 |
| Omics 1 | Latent 1* | Latent 2* | Latent 3* | Noise | Latent 5* | Latent 8 | Latent 9 | Latent 10 | Noise | Noise |
| Omics 2 | Latent 1* | Latent 2* | Noise | Latent 4* | Latent 6* | Latent 8 | Latent 9 | Noise | Latent 11 | Noise |
| Omics 3 | Latent 1* | Noise | Latent 3* | Latent 4* | Latent 7* | Latent 8 | Noise | Latent 10 | Latent 11 | Noise |
| Phenotype Data Latent Factor Composition |  |  |  |  |  |  |  |  |  |  |
| Phenotype | Latent 1*; Latent 2*; Latent 3*; Latent 4*; Latent 5*; Latent 6*; Latent 7* |  |  |  |  |  |  |  |  |  |
|  | 1:20 | 21:40 | 41:60 | 61:80 | 81:100 | 101:160 | 161:220 | 221:280 | 281:340 | 341:1000 |
| Omics 1 | Latent 1* | Latent 2* | Latent 3* | Noise | Latent 5* | Latent 8 | Latent 9 | Latent 10 | Noise | Noise |
| Omics 2 | Latent 1* | Latent 2* | Noise | Latent 4* | Latent 6* | Latent 8 | Latent 9 | Noise | Latent 11 | Noise |
| Omics 3 | Latent 1* | Noise | Latent 3* | Latent 4* | Latent 7* | Latent 8 | Noise | Latent 10 | Latent 11 | Noise |
| Phenotype Data Latent Factor Composition |  |  |  |  |  |  |  |  |  |  |
| Phenotype | Latent 1* |  |  |  |  |  |  |  |  |  |
|  | 1:20 | 21:40 | 41:60 | 61:80 | 81:100 | 101:160 | 161:220 | 221:280 | 281:340 | 341:1000 |
| Omics 1 | Latent 1* | Latent 2* | Latent 3* | Noise | Latent 5* | Latent 8 | Latent 9 | Latent 10 | Noise | Noise |
| Omics 2 | Latent 1* | Latent 2* | Noise | Latent 4* | Latent 6* | Latent 8 | Latent 9 | Noise | Latent 11 | Noise |
| Omics 3 | Latent 1* | Noise | Latent 3* | Latent 4* | Latent 7* | Latent 8 | Noise | Latent 10 | Latent 11 | Noise |
| Phenotype Data Latent Factor Composition |  |  |  |  |  |  |  |  |  |  |
| Phenotype | Latent 2*; Latent 3*; Latent 4*; Latent 5*; Latent 6*; Latent 7* |  |  |  |  |  |  |  |  |  |

The positive “signal” features, intended for model identification, are defined as four-way correlations involving all omics and phenotype, multiple three-way correlations among different combinations of omics with phenotype, and pairwise correlations between each omics and phenotype. Additionally, the nonphenotype-specific correlations are negative “random noise” features to challenge signal feature detection. These include a 3-way correlation among all omics and pairwise correlations among the omics, and these correlation structures serve as noise features, which models should ideally overlook. The performance is evaluated at the node level with Area Under the Receiver Operating Characteristic Curve (AUC) score. A node is predicted positive (signal) if its maximal connection to other nodes in the adjacency matrix passes a certain threshold, which is consistent with the SmCCNet evaluation [24]. The AUC score is then calculated checking prediction results across a series of the threshold value.

We compare the performance of SGTCCA-Net to other multi-omics network analysis methods with 25 replications: (1) STCCA-Net with only the higher order 4-way correlation structure;(2) Best SmCCNet AUC score from various combinations of scaling factors and sparsity levels; (3) Best DIABLO AUC score with combinations of different levels of sparsity and scaling factors. The parameter setup for SmCCNet and DIABLO resulted in a total of 9 models for each method. To ensure a fair comparison between methods, we extracted only one set of canonical weights from each method. More details on parameter settings can be found in S6 Text.

### 0.12 Multi-Omics Data Analysis

In addition to the simulations, we compared the performance of SGTCCA and SmCCA on two data sets. To maintain a fair comparison, we kept all subsequent network analysis steps constant, and therefore did not use DIABLO as the network inference for that method implements a different pipeline. Therefore, we opt for a modified version of SmCCNet (SmCCNet 2.0) [42], ensuring that the network analysis steps align with those of SGTCCA-Net. The best scaling factors and penalty terms selection for SmCCNet are based on 5-fold cross-validation. To be specific, we let the penalty term for each molecular profile vary between 0.25, 0.5, and 0.75, and the scaling factor associated with each phenotype-specific pairwise correlation component specific to the phenotype varies between 1, 5, and 10, and we evaluate the scaled prediction error for each particular combination. In an example of two omics data, if (*a*_1,2_, *b*_1_, *b*_2_) are the scaling factors where *b*_1_ and *b*_2_ represent the phenotype-specific scaling factors and *a*_1,2_ stands for the scaling factor for between-omics correlation, we will scale the scaling factor by *δ* to ensure *a*_1,2_ + *b*_1_ + *b*_2_ = 1, then the best scaling factors and their associated penalty terms are determined by minimizing:

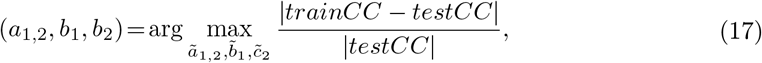

where *trainCC* and *testCC* stand for total training canonical correlation and total testing canonical correlation. All other steps in the pipeline, including network construction and pruning, were kept unchanged. Specifically, we fix steps (2) through (4) of Algorithm 1 and vary the setup for step (1) between SGTCCA and SmCCNet. The study aimed to compare the performance of SGTCCA to SmCCA in extracting the network associated with the phenotype of interest.

#### 0.12.1 TCGA Breast Cancer Network Analysis

The Cancer Genome Atlas Program (TCGA) breast invasive carcinoma project includes RNA sequencing data with normalized counts obtained through the Illumina HiSeq platform, microRNA (miRNA) expression data of tumor samples obtained through the Illumina HiSeq platform at the miRgene-level, and log-ratio normalized reverse phase protein arrays (RPPA) expression data from tumor samples at the gene level. In this experiment, we use tumor purity as the phenotype, defined as the percentage of cancer cells in a sample of tumor tissue [43]. After matching subjects with all available molecular profiles and phenotype data, we obtained a cohort of 105 subjects. Because of their relatively large number, we filtered genes based on the standard deviation to eliminate genes that exhibit lower variability and only include the top 25% of the genes, resulting in a total of 5039 genes, in addition to the 823 miRNAs, and 175 RPPAs. In the data preprocessing step, we regress out age, race, and whether patients received radiation therapy for each of the molecular features to adjust for covariates.

Our SGTCCA-Net pipeline assumes the correlation structure of gene-miRNA-RPPA-phenotype, gene-miRNA-phenotype, gene-RPPA-phenotype, miRNA-RPPA-phenotype, and all pairwise molecular profiles with phenotype. Same as the simulation study, we set the percentage of common subsampled molecular features at 8% and the percentage of distinct molecular features to 2% with a total of 10 iterations when bias subsampling is performed to guarantee sparsity. We use the network pruning algorithm to prune the multi-omics network and set the minimally/maximally final subnetwork size to 30/300.

To evaluate the efficacy of each method, we obtained the NetSHy summarization score for every subnetwork, and correlate the scores with tumor purity. In addition, we calculated the correlation of individual molecular features with tumor purity, contrasting the number of strongly and weakly correlated features between both methods. Since one of the advantages of SGTCCA-Net is its ability to extract higher-order correlation among molecular features, we rank all potential molecular feature triplets and pairs in relation to tumor purity based on their respective higher-order correlation values. This is done to investigate how different molecular features simultaneously interact with respect to tumor purity.

An enrichment analysis was also performed on the aggregate set of genes and proteins through Metascape (v. 3.5.20230501) [44], with the reference set of all genes and protein target genes fed into the SGTCCA-Net pipeline and recognized by Cytoscape (4540). While the aforementioned approach focused solely on genes and proteins, we also extended the enrichment analysis to miRNAs. Using MultiMiR [45], we first identified validated target genes of miRNAs in the subnetwork, then treated target genes that have a non-zero canonical weight in the global network as the enriched set. Enrichment analysis is also performed in Metascape with the same background set as described above.

#### 0.12.2 COPDGene Network Analysis

Phase II COPDGene [46] includes transcriptomics (RNA-Seq), proteomics (SomaLogic 1.3k), and metabolomics (Metabolon) data [47] for studying chronic obstructive pulmonary disease (COPD). For demonstration, we use a smaller cohort to compare different methods. We construct multi-omics networks with respect to forced expiratory volume at one second (FEV_1_). To ensure that we have roughly 1000 genes to run the model, we use standard deviation to filter out genes with less variability (sd < 0.435). After filtering subjects with available data for all molecular profiles and the phenotype, there were 461 subjects with available data for all molecular profiles, with 972 genes, 1305 proteins, and 995 metabolites. To adjust for covariates, we regressed out effects from sex, age, and clinical center for each molecular feature.

Our SGTCCA-Net pipeline assumes the correlation structure of gene-protein-metabolite-phenotype, gene-protein-phenotype, gene-metabolite-phenotype, protein-metabolite-phenotype and all pairwise molecular profiles with phenotype. As in the simulation study and the TCGA breast cancer study, we set the percentage of common subsampled molecular features at 8% and the percentage of distinct molecular features to 2% with a total of 10 iterations when biased subsampling was performed to guarantee sparsity.

To evaluate the efficacy of each method, we obtained the NetSHy summarization scores for every subnetwork, correlating them with FEV_1_. In addition, we calculated the correlation of individual molecular features with FEV_1_, contrasting the number of strongly and weakly correlated features between both methods. Since one of the advantages of SGTCCA-Net is its ability to extract higher-order correlation between molecular features, we rank all potential molecular feature triplets and pairs in relation to FEV1 based on their respective higher-order correlation values. This is done to investigate how different molecular features simultaneously interact with respect to FEV_1_.

Enrichment analysis was also performed on the union set of genes and proteins in the subnetwork through Metascape (v. 3.5.20230501) [44], with the background set to be all genes and protein target genes fed into the SGTCCA-Net pipeline and recognized by Cytoscape (1750). While the aforementioned approach focused solely on genes and proteins, we also extended the analysis to metabolites by running metabolite enrichment analysis on IMPaLa [48] based on the HMDB names of the metabolites in the subnetwork, with the background set to be all metabolites with available HMDB name (227).

FEV1_1_ is a measurement in liters for each subject, which did not take into account body habitus and race. Therefore, we also ran SGTCCA-Net on the FEV1 percent predicted (FEV1_1_PP) to check if the top molecular features identified from FEV1_1_ are also identified in the FEV1_1_PP network, and if the correlation changes. Additionally, we also ran an enrichment analysis in Metascape to check if the pathways overlapped between the two networks.

## Results

### 0.13 Simulation

SGTCCA-Net significantly outperforms the other three methods (SmCCNet, STCCA-Net, and DIABLO) in all simulation study settings and simulated correlation structure designs (Table 2), even when SmCCNet and DIABLO use the parameter setup that performs best in each iteration. In case 1, where all types of phenotype-specific correlation structures are presented and in case 3, when the 4-way correlation structure is removed, the performance of SGTCCA-Net improves by around 10% compared to the best-performing results from SmCCNet and DIABLO, and the level of improvement increases as the number of subjects increases. SGTCCA-Net achieves high signal feature identification accuracy even with only 100 subjects in the presence and absence of different phenotype-specific correlation structures and provides nearly-perfect prediction when the number of subjects doubles, outperforming the other methods by around 15% to 20% in AUC. In case 2, when there is only one type of phenotype-specific correlation structure (4-way correlation), SGTCCA-Net, STCCA-Net, and best-performing DIABLO perform equally well with perfect or near-perfect AUC scores.

**Table 2.** Simulation results. Performance is evaluated through the AUC of the precision-recall curve generated by applying different thresholds to the maximal connection of molecular features to each other. For this simulation, 20 replications are the AUC median and interquartile range in parenthesis of is reported. “Best” AUC for SmCCNet and DIABLO denotes that in each replication, 9 SmCCNet/DIABLO models are run and only the highest AUC score is recorded. The first table is the simulation study for setting 1, which uses latent factors simulated from multivariate normal distribution; the second panel is the simulation study for setting 2, where latent factors are simulated with a highly right-skewed distribution; the third panel is the simulation study for setting 3, where latent factors are simulated with a multivariate normal distribution (same as case 1), but strong random noise is enforced on omics data. Each simulation study contains 3 cases: Case 1 means that the signal molecular features are defined as all 4-way, 3-way, and pairwise phenotype-specific correlation structure; case 2 removes the phenotype-specific 4-way correlation structure; case 3 removes the phenotype-specific 3-way and pairwise correlation structure. In each case, the data is simulated with 100 or 200 subjects.

| Normal Latent Factor: Median AUC (Interquartile Range) |  |  |  |  |  |
| --- | --- | --- | --- | --- | --- |
| Method | n | SGTCCA-Net | STCCA-Net | SmCCNet (Best) | DIABLO (Best) |
| Case 1 | n = 100 | 0.885 (0.859, 0.906) | 0.656 (0.604, 0.680) | 0.706 (0.689, 0.736) | 0.745 (0.718, 0.780) |
|  | n = 200 | 0.978 (0.966, 0.986) | 0.694 (0.614, 0.738) | 0.742 (0.720, 0.818) | 0.825 (0.754, 0.880) |
| Case 2 | n = 100 | 1.000 (1.000, 1.000) | 0.995 (0.955, 1.000) | 0.825 (0.779, 0.853) | 0.931 (0.920, 0.954) |
|  | n = 200 | 1.000 (1.000, 1.000) | 1.000 (1.000, 1.000) | 0.823 (0.811, 0.837) | 0.939 (0.931, 1.000) |
| Case 3 | n = 100 | 0.890 (0.864, 0.942) | 0.622 (0.583, 0.701) | 0.710 (0.679, 0.740) | 0.701 (0.670, 0.757) |
|  | n = 200 | 0.977 (0.965, 0.983) | 0.679 (0.635, 0.711) | 0.766 (0.726, 0.816) | 0.814 (0.786, 0.891) |

| Skewed Latent Factor: Median AUC (Interquartile Range) |  |  |  |  |  |
| --- | --- | --- | --- | --- | --- |
| Method | n | SGTCCA-Net | STCCA-Net | SmCCNet (Best) | DIABLO (Best) |
| Case 1 | n = 100 | 0.845 (0.814, 0.884) | 0.686 (0.603, 0.730) | 0.705 (0.664, 0.722) | 0.713 (0.675, 0.768) |
|  | n = 200 | 0.935 (0.921, 0.954) | 0.729 (0.675, 0.788) | 0.756 (0.726, 0.804) | 0.847 (0.813, 0.887) |
| Case 2 | n = 100 | 1.000 (1.000, 1.000) | 1.000 (0.980, 1.000) | 0.824 (0.780, 0.846) | 0.939 (0.932, 1.000) |
|  | n = 200 | 1.000 (1.000, 1.000) | 1.000 (1.000, 1.000) | 0.817 (0.770, 0.855) | 0.949 (0.933, 1.000) |
| Case 3 | n = 100 | 0.855 (0.799, 0.891) | 0.677 (0.614, 0.719) | 0.714 (0.681, 0.748) | 0.725 (0.651, 0.777) |
|  | n = 200 | 0.936 (0.925, 0.949) | 0.755 (0.678, 0.795) | 0.712 (0.687, 0.735) | 0.803 (0.738, 0.839) |

**Table 2.** Simulation results. Performance is evaluated through the AUC of the precision-recall curve generated by applying different thresholds to the maximal connection of molecular features to each other.
| Noisy Omics Data: Median AUC (Interquartile Range) |  |  |  |  |  |
| --- | --- | --- | --- | --- | --- |
| Method | n | SGTCCA-Net | STCCA-Net | SmCCNet (Best) | DIABLO (Best) |
| Case 1 | n = 100 | 0.776 (0.761, 0.805) | 0.578 (0.554, 0.595) | 0.702 (0.677, 0.724) | 0.703 (0.667, 0.780) |
|  | n = 200 | 0.909 (0.892, 0.919) | 0.603 (0.579, 0.638) | 0.785 (0.748, 0.801) | 0.813 (0.766, 0.920) |
| Case 2 | n = 100 | 0.998 (0.996, 1.000) | 0.777 (0.666, 0.880) | 0.839 (0.789, 0.853) | 1.000 (0.918, 1.000) |
|  | n = 200 | 1.000 (1.000, 1.000) | 0.921 (0.822, 0.959) | 0.832 (0.815, 0.846) | 1.000 (0.936, 1.000) |
| Case 3 | n = 100 | 0.781 (0.746, 0.809) | 0.556 (0.538, 0.614) | 0.710 (0.678, 0.729) | 0.701 (0.670, 0.757) |
|  | n = 200 | 0.916 (0.888, 0.936) | 0.583 (0.565, 0.609) | 0.782 (0.745, 0.798) | 0.799 (0.778, 0.876) |

When the underlying latent factors are highly right-skewed and omics data are noisy, the performance of all models are impacted, but SGTCCA-Net still outperforms the other methods. This indicates that even when the signal is masked by noise to some extent, or normality is not guaranteed, SGTCCA-Net still performs well compared to other methods, and its performance improves as the sample size increases.

### 0.14 TCGA Breast Cancer Network Analysis

After running a network analysis pipeline with SGTCCA and SmCCNet for the phenotype of tumor purity, the final network size for both SGTCCA-Net and SmCCNet is 300. Specifically, there are 128 genes, 125 miRNAs, and 47 proteins in the SGTCCA network, while there are 128 genes, 59 miRNAs, and 113 proteins in the SmCCNet network. The NetSHy network summarization score correlation with respect to tumor purity is 0.689 and 0.500 for SGTCCA-Net and SmCCNet respectively. In the SGTCCA-Net network, there are 55 molecular features with weak correlation with respect to tumor purity (FDR-adjusted > 0.05), while for SmCCNet there are 105 such molecular features. To check whether each method includes these nodes incorrectly or whether they contribute to the network, we ran the NetSHy network summarization score on these weak nodes only, extracted the first 10 PCs, and calculated the maximum correlation between the PC score and tumor purity. For SGTCCA-Net, the maximum correlation is -0.232 (*p* = 0.017), while for SmCCNet, the maximum correlation is only -0.176 (*p* = 0.072). This implies that weak nodes within the SGTCCA-Net network are more significantly associated with tumor purity compared to those in the SmCCNet network. Furthermore, a total of 139 molecular features are shared between the SGTCCA-Net and SmCCNet methods, with each method having 161 unique molecular features. Of the 161 unique molecular features in each network, 125 and 66 are significantly associated with tumor purity (FDR < 0.05) for SGTCCA-Net and SmCCNet, respectively.

We calculate the PageRank score for the SGTCCA-Net subnetwork and present the top 5 nodes for each molecular profile in Table 3. We observed that all of the molecular features in the table are significantly correlated with tumor purity. *ACVRL1* and *KAT2A* are the only two nodes with low correlation to tumor purity (FDR > 0.001). To check whether these nodes contribute to the network, we examine the correlation between these two nodes and other molecular features of the subnetwork. We observed that *ACVRL1* is highly correlated with *PCNA* (*ρ*_*between*−*omics*_ = 0.474), which is highly correlated with tumor purity (*ρ*_*omics*−*pheno*_ = −0.564) and *KAT2A* (*ρ*_*between*−*omics*_ = 0.476), which is highly correlated with tumor purity (*ρ*_*omics*−*pheno*_ = −0.564).

**Table 3.** Top 5 molecular features from each molecular profile and their individual correlation with respect to tumor purity for TCGA breast cancer data (with p-value).

| Type | Name | Correlation | FDR |
| --- | --- | --- | --- |
| Gene (128) | TMC8 | -0.712 | <0.001 |
|  | ARHGAP9 | -0.752 | <0.001 |
|  | SPIB | -0.638 | <0.001 |
|  | HCST | -0.712 | <0.001 |
|  | IRF8 | -0.748 | <0.001 |
| miRNA (125) | hsa-mir-142 | -0.564 | <0.001 |
|  | hsa-mir-146a | -0.560 | <0.001 |
|  | hsa-mir-150 | -0.593 | <0.001 |
|  | hsa-mir-155 | -0.534 | <0.001 |
|  | hsa-let-7i | -0.572 | <0.001 |
| Protein (47) | PCNA | -0.564 | <0.001 |
|  | CCNB1 | -0.409 | <0.001 |
|  | NRG1 | -0.466 | <0.001 |
|  | KAT2A | 0.324 | 0.001 |
|  | ACVRL1 | 0.297 | 0.003 |

The top 5 ontologies from the enrichment analysis for the SGTCCA-Net final subnetwork (Figure 3) includes lymphocyte activation (49/186 genes enriched), positive regulation of immune response (45/204 genes enriched), adaptive immune response (38/145 genes enriched), regulation of leukocyte activation (46/227 genes enriched), and Adaptive Immune System (37/178 genes enriched). Compared to enrichment analysis result from SmCCNet, we observe that the ontology results from both networks fall into similar categories. However, when only looking at the immune response-related ontologies, a stronger enrichment analysis result is identified in SGTCCA-Net network (S1 Fig). Additionally, using MCODE, we observed 6 modules of the protein-protein interaction network as shown in Fig 3b. For example, in the first network module (red), one of the proteins is *CD3D*, which has prognostic potential for breast cancer and associated with lymphocyte infiltration and immune checkpoints [49].

**Fig 3.**
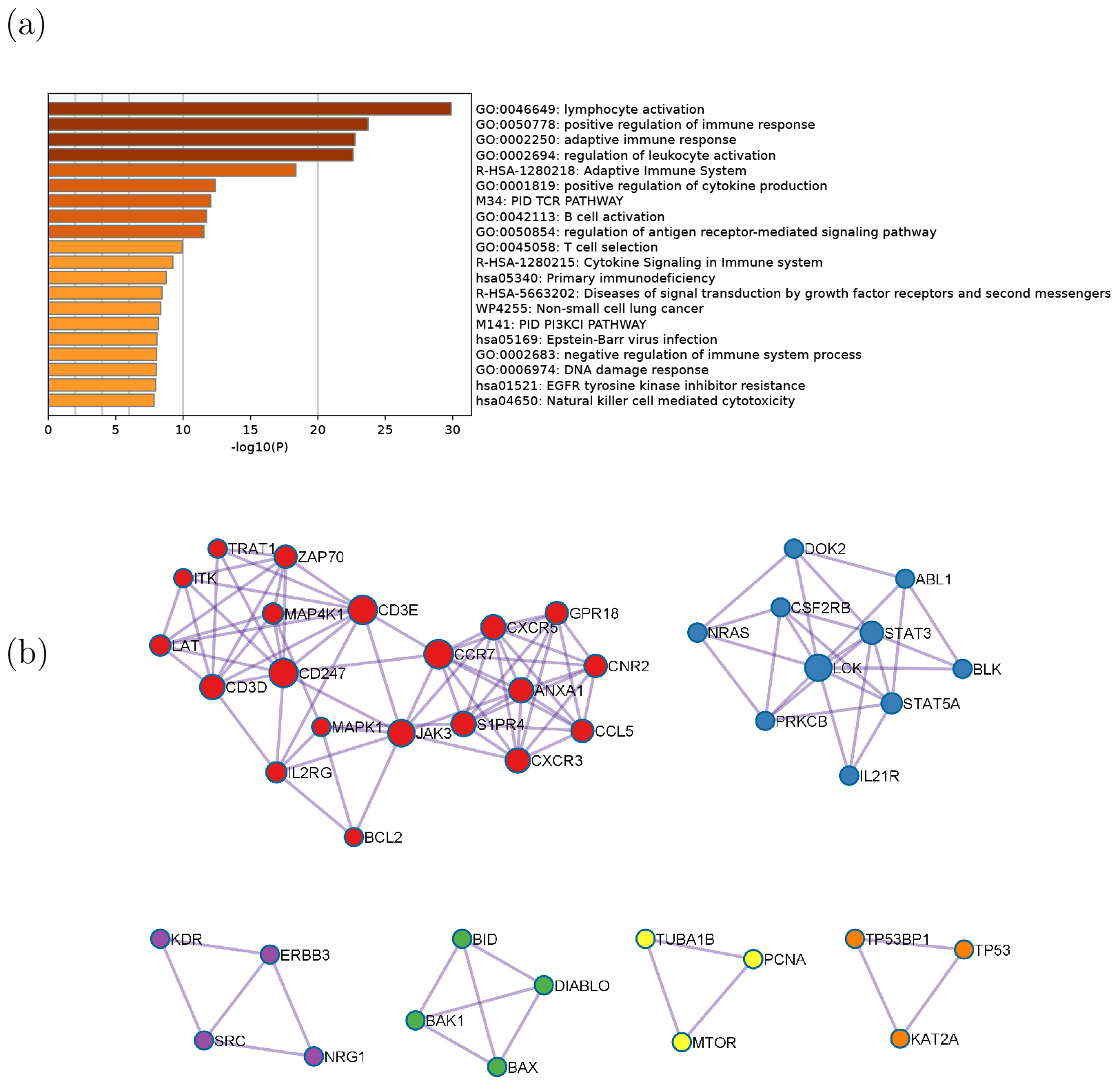
Enrichment analysis results for TCGA breast cancer data with respect to tumor purity. (a) The top pathways that are associated with the final network. (b) Protein-protein interaction (PPI) network for the multi-omics network from SGTCCA-Net with respect to tumor purity colored by clusters. Clusters are generated based on the Molecular Complex Detection (MCODE) algorithm.

We ran MultiMiR on the miRNAs from each subnetwork. For the SGTCCA-Net network, among 1288 global network genes with non-zero canonical weight, 179/697 of them are the validated/predicted target genes of miRNAs in the subnetwork; while for the SmCCNet network, among 1024 global network genes with non-zero canonical weight, only 102/212 of them are the validated/predicted target genes of miRNAs in the subnetwork. This result suggests that more connections between genes and miRNAs are found in the SGTCCA-Net network compared to the SmCCNet network. The enrichment analysis based on the 179 validated targets of SGTCCA-Net subnetworks (S2 Fig) shows that the ontologies fall into similar categories compared to the gene/protein enrichment analysis, including lymphocyte differentiation and adaptive immune system.

We examined the top combinations (4/3-way correlation) and found that the top 4-way correlation implies the top 3-way correlation in some cases (S2 Table). For example, it was observed that *hsa-mir-3133* and *KDR* dominate the 4-way relationship between gene, miRNA, protein, and tumor purity, which also implies the 3-way correlation between *hsa-mir-3133, KDR*, and tumor purity. Interestingly, *BEND4*, one of the genes present in the top 4-way correlation, is shown to be the predicted target of *hsa-mir-3133* with a predicted score of 0.728, and *KDR* is also the predicted target of *hsa-mir-3133* with a predicted score of 0.572. Through examining the 3-way correlation we can find that even though the top 3-way correlation could imply the 4-way correlation, the signal is generally weaker. For instance, it was found that *hsa-mir-142* dominates the gene-miRNA-tumor purity correlation, *PCNA* dominates both the gene-protein-tumor purity correlation and the miRNA-protein-tumor purity correlation, and *CCNB1* dominates the miRNA-protein-tumor purity correlation (details in S2 Table). However, it was observed that the highest 4-way correlation involved in *hsa-mir-142* is 2.405 (rank 94/631,680 across all 4-way correlation), in *PCNA* is 2.615 (rank 59/631,680 across all 4-way correlation), in *CCNB1* is 2.093 (rank 191/631,680 across all 4-way correlation). A particular example is the 3-way correlation that involves *CCNB1* and *hsa-mir-150*, which is observed in the top 10 3-way correlation between miRNAs, proteins and tumor purity, but the highest 4-way correlation involves these two molecular features is only 1.914 (rank 398/631,680). This suggests that some 3-way correlation combinations may be ignored by the 4-way correlation, and indicates the importance of incorporating the 3-way correlation into the model.

Lastly, the PageRank score for each molecular feature in the final subnetwork was calculated, and the top 10 molecular features from each molecular profile were selected for network visualization in Figure 4. It was noticed that higher connectivity occurs in proteins *PCNA* and *CCNB1* (both appear in Table 3 and S2 Table), and both of them are observed in the top 3-way miRNA-protein-tumor purity correlation and found as prognostic biomarkers for breast cancer [50, 51].

**Fig 4.**
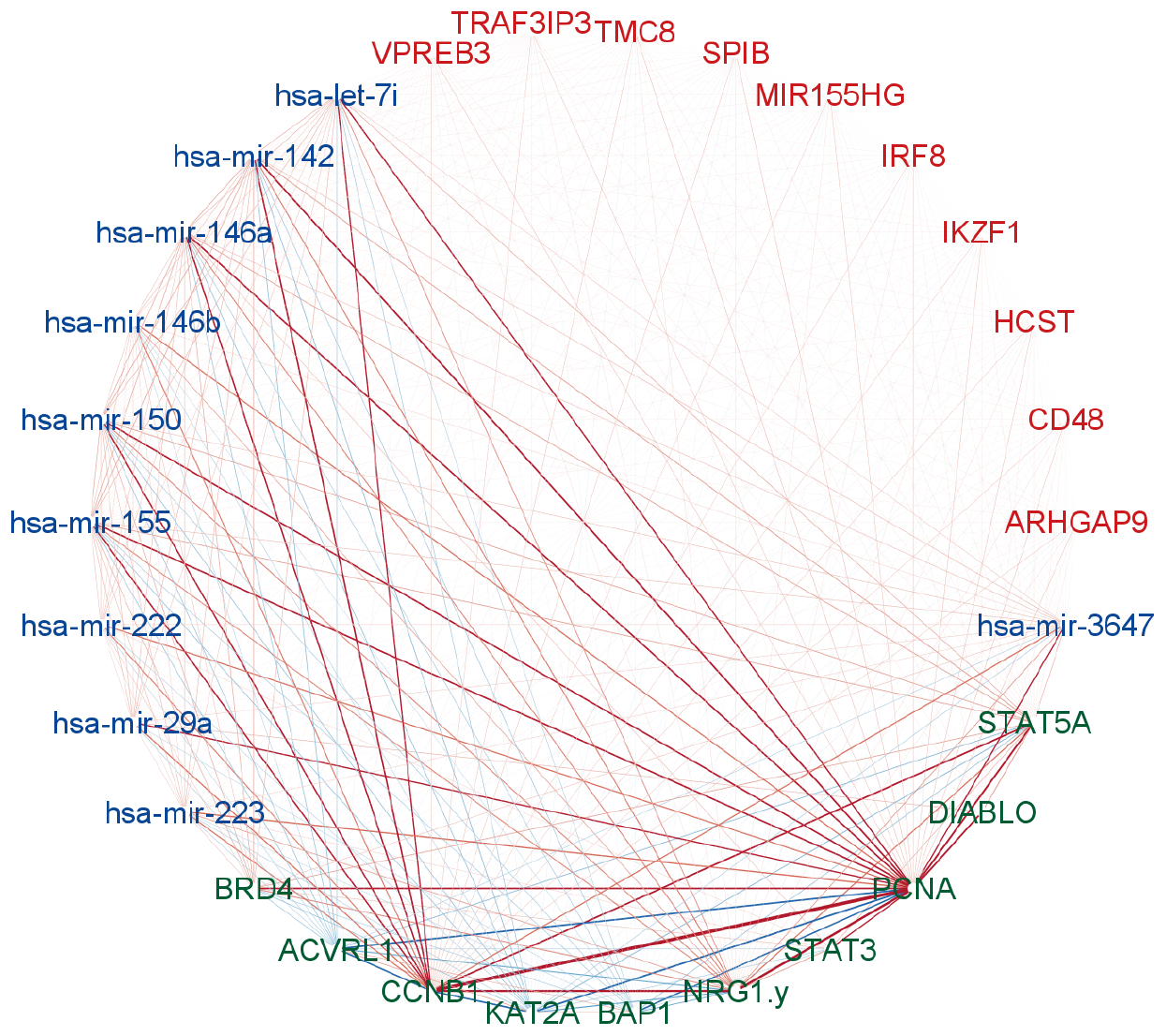
SGTCCA-Net network with top 10 molecular features from each molecular profile. Multi-omics network module for TCGA breast cancer data with respect to tumor purity. Nodes are genes (red), miRNAs (blue) and RPPAs (green). The edge color denotes positive correlation (red) or negative correlation (blue) between molecular features with the width denoting the strength of the connection. Edges are filtered based on Pearson correlation with a threshold of 0.2.

### 0.15 COPDGene Network Analysis

After running a network analysis pipeline with SGTCCA-Net and SmCCNet for the FEV1 phenotype, the final network size for SGTCCA-Net is 170, while SmCCNet has a final network size of 117. The NetSHy summarization score for SGTCCA-Net is correlated with FEV_1_ with a correlation of 0.211, while for SmCCNet it is only 0.162. In the SGTCCA-Net network, there are 58 molecular features with weak correlation with respect to FEV_1_ (FDR > 0.05), while for SmCCNet, there are 82 such molecular features. To check whether each method includes these nodes incorrectly or whether they contribute to the network, we extracted the NetSHy summarization score on these weak nodes only, extracted the first 3 PCs, and calculated the maximum correlation between PC score and FEV1. For SGTCCA-Net, the maximum correlation is -0.159 (*p <* 10^−3^), while for SmCCNet, the maximum correlation is only 0.090 (*p* = 0.053). This implies that the weak nodes included in SGTCCA-Net are shown to be more related to the phenotype than the weak nodes included in SmCCNet. Furthermore, a total of 29 molecular features are shared between the SGTCCA-Net and SmCCNet methods, with each method having 141 and 88 distinct molecular features respectively. Of the 141 unique molecular features in the SGTCCA-Net network, 89 of them are significantly associated with FEV_1_ (FDR < 0.05), while of 88 unique molecular features in the SmCCNet network, only 14 of them are significantly associated with FEV1 (FDR < 0.05). In particular, *Troponin T* (*ρ* = −0.278, *p <* 10^−3^) and *C-reactive protein* (*ρ* = −0.190, *p <* 10^−3^) have a strong correlation with respect to FEV1 and are not identified by SmCCNet, and studies have shown that elevated *Troponin T* levels during exacerbation are associated with death in patients with COPD [52, 53], and more validation will need to be conducted to evaluate whether it is related to FEV_1_ at the stable state.

We calculate the PageRank score for the SGTCCA-Net subnetwork and present the top 5 nodes for each molecular profile in Table 4. We observed that most of the molecular features in the table are significantly correlated with FEV1 (FDR < 0.05), with exceptions such as *Neutrophil gelatinase-associated lipocalin* and *1-stearoyl-2-oleoyl-GPI (18:0/18:1)\**. To check whether these nodes contribute to the network, we examined the correlation between these two nodes and other molecular features of the subnetwork and observed that *Neutrophil gelatinase-associated lipocalin* is highly correlated with *N-acetylneuraminate* (*ρ*_*between*−*omics*_ = 0.382), which is highly correlated to FEV1 (*ρ*_*omics*−*pheno*_ = −0.134) and *1-stearoyl-2-oleoyl-GPI (18:0/18:1)\** is highly correlated with s *1-(1-enyl-stearoyl)-GPE (P-18:0)\** (*ρ*_*between*−*omics*_ = 0.509), which is highly correlated to FEV1 (*ρ*_*omics*−*pheno*_ = 0.151).

**Table 4.**
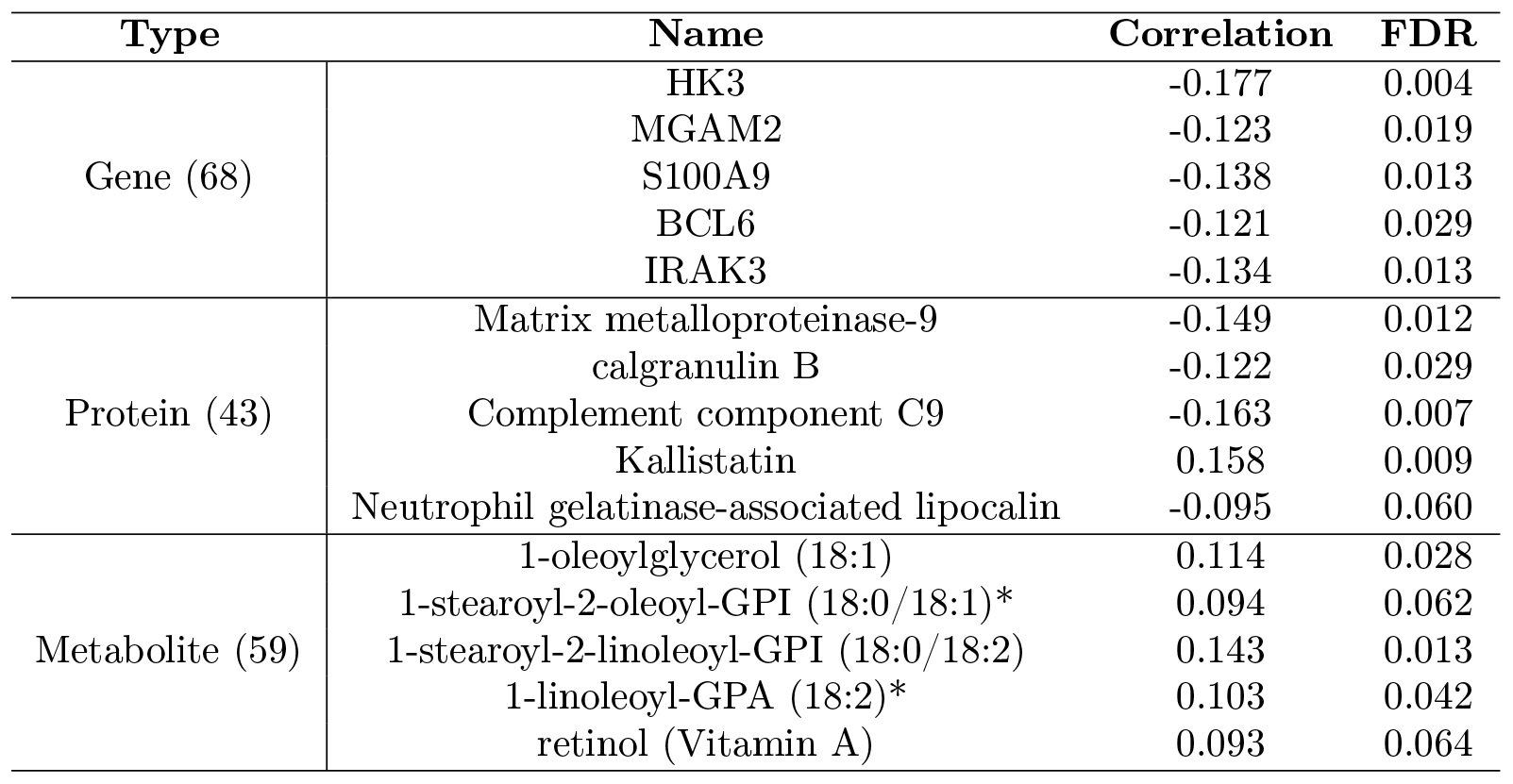
Top 5 molecular features from each molecular profile for SGTCCA-Net based on the PageRank and their individual correlation with respect to FEV1 for COPDGene data (with p-value).

Pathways such as regulation of TLR by endogenous ligand (7/13 genes enriched) and neutrophil degranulation (23/102 genes enriched) are strongly associated with the severity of COPD from the enrichment analysis (Figure 5a). Toll-like receptor (TLR) 2 is elevated in monocytes from individuals with COPD [54], and increased degranulation in COPD is found to increase on the surface of unstimulated neutrophils in patients with COPD and can cause additional airway damage in patients with COPD [55]. In particular, when checking the correlation between the network and cell type, we found that NetSHy PC1 is highly correlated with the percentage of neutrophil cells (*ρ* = −0.767), which may explain the significance of the neutrophil degranulation pathway. However, we did not regress out cell type composition in advance as they can reflect disease state.

**Fig 5.**
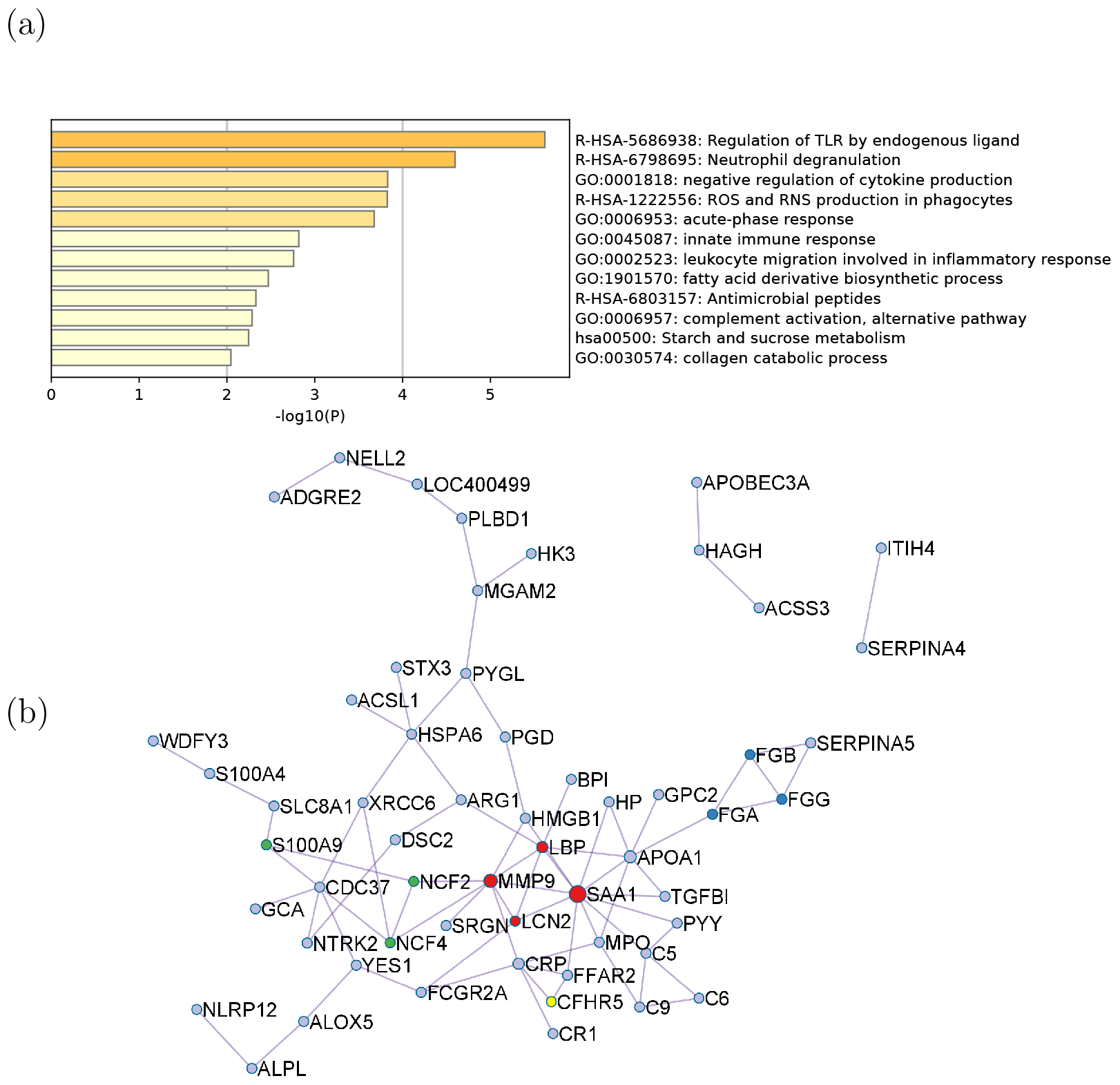
Enrichment analysis results for COPDGene data with respect to FEV1. (a) The top pathways that are associated with the final network. (b) Protein-protein interaction (PPI) network for the multi-omics network from SGTCCA-Net with respect to FEV1 colored by clusters. Clusters are generated based on the Molecular Complex Detection (MCODE) algorithm.

We ran the enrichment analysis on the metabolites of the SGTCCA-Net subnetwork and found that most of the top pathways are associated with phospholipid/glyerophospholipid and bile acid (S4 Table). Compared to non-smokers, current smokers have decreased phospholipid levels regardless of COPD status [56].

Figure 5b shows that there are three groups of protein-protein interaction based on MCODE, and the largest group is centered on the group with SAA1 and MMP9 (red) with the enrichment score of 1.5. SAA1 is related to the acute exacerbation of chronic obstructive pulmonary disease [57], and higher levels of MMP-9 correspond to a higher influx of neutrophils and lymphocytes, signaling an exacerbation of COPD in which a higher burden of MMP-9 is observed in the airways [58].

We examined the top combinations (4/3-way correlation) and found that the top 4-way correlation implies the top 3-way correlation in some cases (S3 Table). For instance, it was observed that *MTCO1P12, Tyrosine-protein kinase Yes*, and *C-glycosyltryptophan* dominate the 4-way relationship between gene, protein, metabolite, and FEV_1_, which also implies the 3-way correlation between *Tyrosine-protein kinase Yes* (the protein product of *YES1*), *C-glycosyltryptophan* and FEV_1_. Through examining the 3-way correlation we can find that even though the top 3-way correlation could imply the 4-way correlation but the signal is generally weaker. For instance, it was found that *Matrix metalloproteinase-9* dominates the gene-protein-fev1 correlation, *1-stearoyl-2-oleoyl-GPI (18:0/18:1)\** dominates the gene-metabolite-fev1 correlation, and *C-glycosyltryptophan* and *iminodiacetate (IDA)* dominate the protein-metabolite-fev1 correlation (details in S3 Table). However, it was observed that the highest 4-way correlation involved in *Matrix metalloproteinase-9* is 0.431 (rank 233/172,516 across all 4-way correlation), in *iminodiacetate (IDA)* is 0.580 (rank 30/172,516 across all 4-way correlation). A particular example is the 3-way correlation that involves *Matrix metalloproteinase-9* and other genes, we found that all top 10 3-way correlation between gene, protein, and FEV_1_ involve this protein, but only 2 of the genes are shown in the top 1000 4-way correlation along with *Matrix metalloproteinase-9*. This suggests that some 3-way correlation combinations may be ignored by the 4-way correlation, and indicates the importance of incorporating the 3-way correlation into the model.

Furthermore, the PageRank score for each molecular feature in the final subnetwork was calculated, and the top 10 molecular features from each molecular profile were selected for visualization of the network in Fig. 6. In particular, the C-reactive protein has been shown to be associated with poor lung function [59]; the expression level of S100A9 is elevated in patients with COPD, which triggers neutrophil degranulation and releases inflammatory and proteolytic enzymes [60].

**Fig 6.**
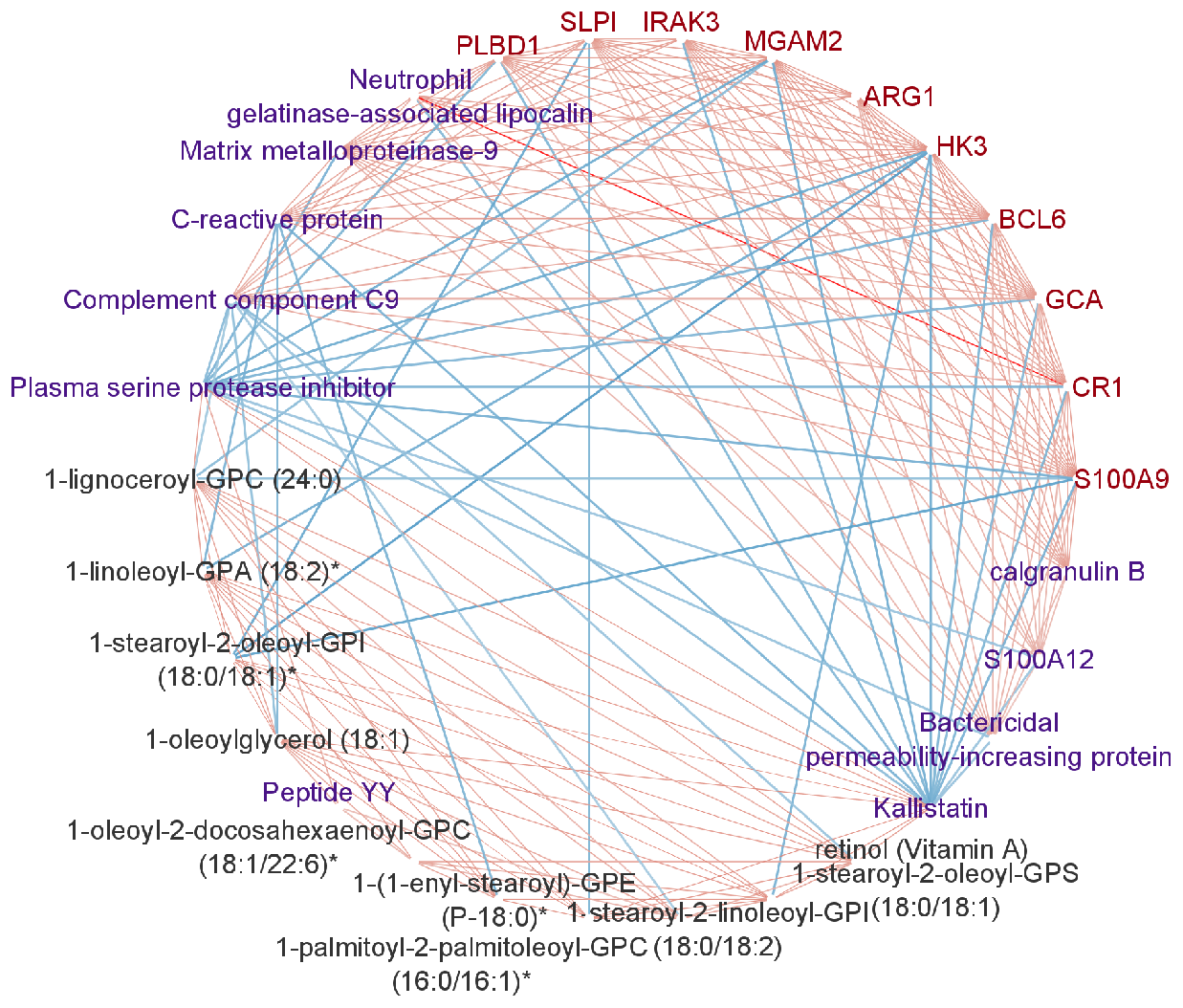
SGTCCA-Net network with top 10 molecular features from each molecular profile. Multi-omics network module for COPDGene data with respect to FEV_1_. The red nodes stand for genes, the purple nodes stand for proteins, and the black nodes stand for metabolites. The width and color depth of the edge stands for the strength of the connection between two molecular features and the type of color stands for whether two nodes are positively correlated (red) or negatively correlated (blue). Edges are filtered based on Pearson correlation with a threshold of 0.2.

We also repeated the analysis on FEV1 percent predicted (FEV1_1_PP), which takes into account body habitus and race. After applying the SGTCCA-Net to FEV1_1_PP, we identified a final subnetwork with 43 features. In particular, 16 of these features exhibit a strong correlation with FEV1_1_PP (with |*ρ*| *>* 0.15), and are all present in the FEV1_1_ final subnetwork. This significant overlap between the FEV1_1_ and FEV1_1_PP subnetworks suggests a consistent molecular pattern that influences both metrics. Furthermore, Metascape enrichment analysis highlighted similar patterns in both FEV1_1_ and FEV1_1_PP networks, with shared pathways including acute-phase response, inflammatory response, and neutrophil degranulation (S3 Fig). Interestingly, these molecular features generally show a higher correlation with FEV1_1_PP than with FEV1_1_ (detailed in S5 Table). This observation demonstrates that the associations between top-correlated molecular features and pulmonary function remain robust, unaffected by variations in body habitus and race.

## Discussion

In this work, we introduce a new multi-omics network analysis pipeline, Sparse Generalized Tensor Canonical Correlation Analysis Network Inference (SGTCCA-Net), designed for biomarker identification through both higher-order and lower-order correlations, thereby creating molecular interaction networks pertinent to specific phenotypes. We propose a new approach, Sparse Generalized Tensor Canonical Correlation Analysis (SGTCCA), which extends Tensor Canonical Correlation Analysis to accommodate multiple correlation structures at once, ensuring sparsity via the biased subsampling technique. Additionally, we bring forth a novel network analysis and network pruning algorithm to derive multi-omics subnetwork modules associated with the phenotype of interest.

Contrary to existing CCA-based multi-omics network inference approaches like SmCCNet and DIABLO, which only examine pairwise correlations, SGTCCA-Net encapsulates both higher and lower-order correlations. Our simulation analyses demonstrate that SGTCCA-Net is adept at pinpointing features with diverse phenotype-specific correlation structures, thereby surpassing the other two CCA-based methods across varied simulation settings.

Our real data analysis highlights the ability of SGTCCA-Net to identify molecular features for diseases such as cancer and chronic obstructive pulmonary disease (COPD). Interestingly, the biomarkers identified are not limited to those with strong phenotype correlations; we also identified molecular features with weak phenotype correlations that are highly connected to other nodes and contribute to the network. Furthermore, we validated our findings using various techniques, including enrichment analysis of biological pathways and miRNA-target relationships. These validations confirm the enhanced performance of SGTCCA-Net over SmCCNet and the ability to find relevant pathways. Furthermore, we also demonstrate the top higher-order correlation associated with the final subnetwork. This study showcases the potential for the integration of multi-omics data to improve our understanding of complex diseases, such as COPD and breast cancer, and highlights the utility of SGTCCA-Net as a powerful tool for analyzing such data.

Beyond its primary use in multi-omics network inference, SGTCCA has broader applications, and can be useful for multi-view dimensional reduction tasks. It can be used to project omics data into a shared lower-dimensional space, suitable for both supervised tasks, such as disease classification, and unsupervised tasks, such as subtyping.

## Supporting information

**S1 Fig. Enrichment analysis ontology results based on the joint set of gene/protein for TCGA breast cancer data and COPDGene data for SmCCNet subnetwork**. The top pathways that are associated with the final network of SmCCNet based on Metascape. a: Top ontology pathways for TCGA breast cancer data with respect to tumor purity; b: Top ontology pathways for COPDGene data with respect to FEV1.

**S2 Fig. Results of the enrichment analysis ontology based on the validated miRNA target genes for the TCGA breast cancer data for the SGTCCA-Net subnetwork**. Top ontology pathways based on validated miRNA target genes for the breast cancer data from TCGA with respect to tumor purity.

**S3 Fig. Enrichment Analysis Results of SGTCCA-Net for** FEV_1_ **Percent Predicted:** This figure focused on pathways enriched in the joint gene/protein set derived from the COPDGene data, specifically within the SGTCCA-Net subnetwork associated with FEV_1_ Percent Predicted. The identified pathways provide insights into the molecular mechanisms linked to pulmonary function as measured by FEV_1_ Percent Predicted.

**S1 Text. Higher-order Covariance Tensor for both Odd and Even Number of Views**

**S2 Text. Proof of Theorem 1**

**S3 Text. Sparse Generalized Tensor Canonical Correlation Analysis (SGTCCA) Algorithm**

**S4 Text. Biased Subsampling Covariance Density**

**S5 Text. Network Analysis and Network Pruning Algorithm S6 Text. Additional Detail of Simulation Study Setup**

**S1 Table. Simulation Setup Table**. Summary table of all simulation scenarios. Starting with 3 omics data simulation settings: (1) normal latent factors, (2) highly right-skewed latent factors, and (3) noisy omics data. Each of these settings has 3 cases: case 1 with all phenotype-specific correlation structures; case 2 with only 4-way phenotype-specific correlation structure; and (3) case 3 with 3-way and pairwise phenotype-specific correlation structure. All these cases will be evaluated based on sample sizes of 100 and 200. Σ represents a diagonal matrix with diagonal values to be 1.

**S2 Table. Top Network Higher-order Correlation for TCGA Breast Cancer Network**. Top 5 network higher-order correlation of breast cancer data of different correlation structures.

**S3 Table. Top network higher-order correlation for the COPDGene network**. Top 5 network higher-order correlation of COPDGene data of different correlation structures.

**S4 Table. Top Metabolite Enrichment Pathways**. Top 10 pathways of metabolite enrichment analysis result from IMPaLa for SGTCCA-Net subnetwork.

**S5 Table. Overlap Molecular Features between FEV1 and FEV1 Percent Predicted Network** Molecular features overlap between FEV1 and FEV1 percent predicted network generated from SGTCCA-Net (where absolute correlation with phenotype >0.15).

## Implementation

The model, simulations, and real data experiment were implemented using R version 4.2.2. All code and scripts used in this study are publicly available via GitHub at https://github.com/liux4283/SparseGTCCANet.

## Availability

The TCGA breast cancer data used in the real data experiment section is available at: http://linkedomics.org/data_download/TCGA-BRCA/. Clinical data and SOMAScan data are available through COPDGene (https://www.ncbi.nlm.nih.gov/gap/, ID: phs000179.v6.p2). RNA-Seq data are available through dbGaP (https://www.ncbi.nlm.nih.gov/gap/, ID: phs000765.v3.p2).

Metabolon data is available at Metabolomics Workbench (https://www.metabolomicsworkbench.org/ ID: PR000907). The source code for SGTCCA-Net is available at https://github.com/liux4283/SparseGTCCANet.

## Acknowledgments

We thank all members of the NetCO team for all the insightful feedback on this novel pipeline. This work is also supported in part by funds from the National Heart, Lung, and Blood Institute, National Institutes of Health (R01 HL152735 and TransOmics for Precision Medicines Fellowship).

This work was also supported by NHLBI grants U01 HL089897 and U01 HL089856 and by NIH contract 75N92023D00011. The COPDGene study (NCT00608764) has also been supported by the COPD Foundation through contributions made to an Industry Advisory Committee that has included AstraZeneca, Bayer Pharmaceuticals, Boehringer-Ingelheim, Genentech, GlaxoSmithKline, Novartis, Pfizer, and Sunovion.

## Notes

https://github.com/liux4283/SparseGTCCANet

### Competing Interest Statement

The authors have declared no competing interest.

