## Supplementary Material for "A Generalized Higher-order Correlation Analysis Framework for Multi-Omics Network Inference"

### S1 Figure: Enrichment Analysis Result from SmCCNet

(a)

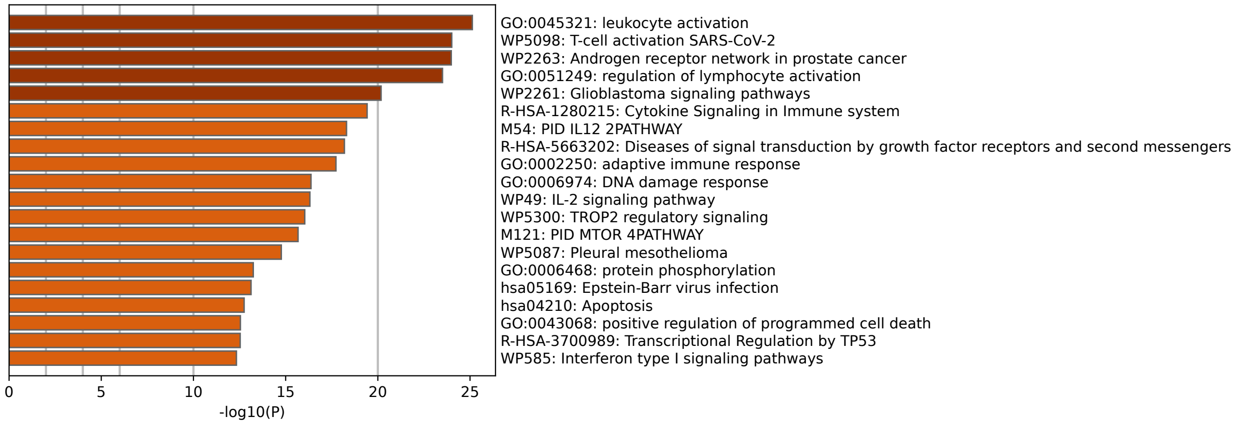


(b)

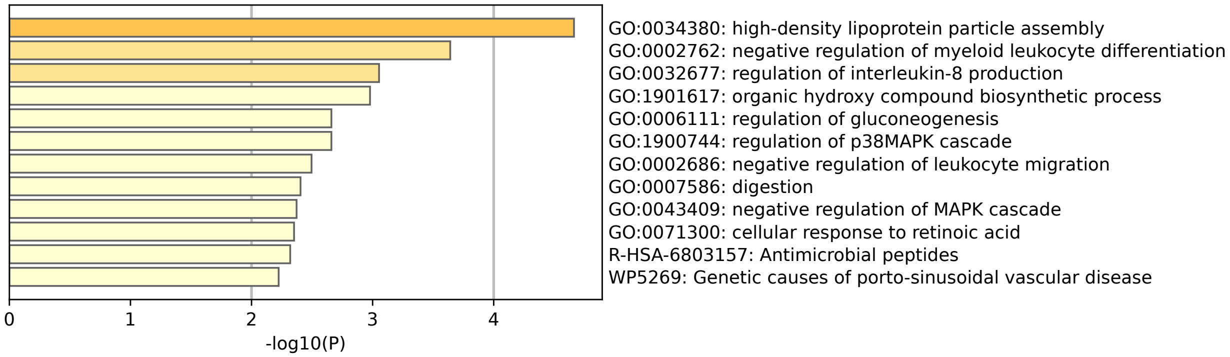


Figure 1: **Enrichment analysis ontology results based on the joint set of gene/protein for TCGA breast cancer data and COPDGene data for SmCCNet subnetwork.** The top pathways that are associated with the final network of SmCCNet based on Metascape. a: Top ontology pathways for TCGA breast cancer data with respect to tumor purity; b: Top ontology pathways for COPDGene data with respect to FEV1.

### S2 Figure: Enrichment Results with Validated Target Genes of MiRNAs

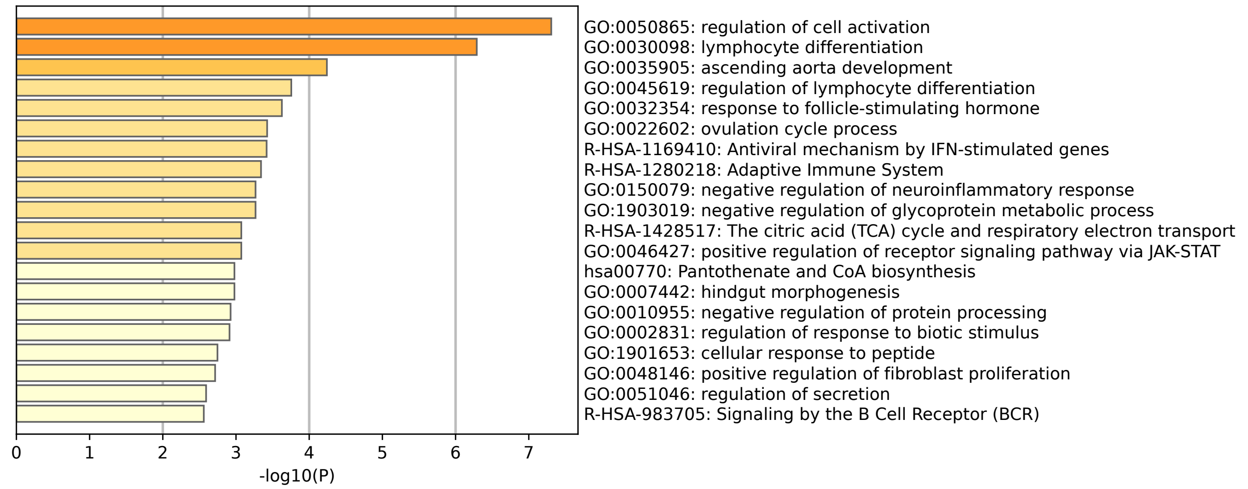


Figure 2: **Enrichment analysis ontology results based on the validated target genes of miRNAs for TCGA breast cancer data for SGTCCA-Net subnetwork.** Top ontology pathways based on validated miRNA-target genes for TCGA breast cancer data with respect to tumor purity.

### S3 Figure: Enrichment Analysis Result from SGTCCA-Net for $\text{FEV}_1$ Percent Predicted

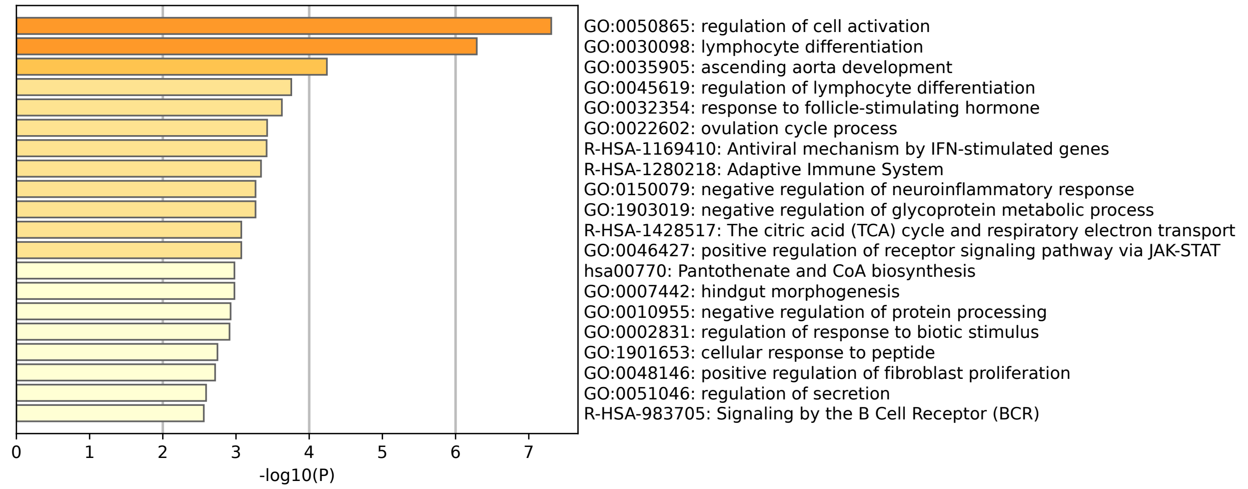


Figure 3: **Enrichment Analysis Results of SGTCCA-Net for** $\text{FEV}_1$ **Percent Predicted:** This figure focused on pathways enriched in the joint gene/protein set derived from the COPDGene data, specifically within the SGTCCA-Net subnetwork associated with $\text{FEV}_1$ Percent Predicted. The identified pathways provide insights into the molecular mechanisms linked to pulmonary function as measured by $\text{FEV}_1$ Percent Predicted.

### S1 Appendix: Higher-order Covariance Tensor for Both Odd and Even Number of Views

Let $z_1, z_2, ..., z_k \in \mathbb{R}^{n\times 1}$ be $k$ vectors that are centered and scaled, then the higher-order covariance between these vectors can be defined as:

$$\rho(z_1, z_2, ..., z_k) = \begin{cases} |\frac{1}{n}(z_1 z_2 ... z_k)^T \mathbf{1}|, \\ \qquad\qquad\qquad\qquad \text{if } k = 2m, m \in \mathbb{N} \\ \frac{1}{n\cdot k}\sum_{j=1}^{k} |(z_1 ... |z_j| ... z_k)^T \mathbf{1}|, \\ \qquad\qquad\qquad\qquad \text{if } k = 2m+1, m \in \mathbb{N}, \end{cases} \tag{1}$$

where $z_1 z_2 ... z_k$ is the element-wise multiplication between $k$ vectors. The reason for calculating the higher-order correlation in different ways for odds and even number of views is that the higher-order correlation calculation for the odd number of views suffers from effect cancellation. For instance, suppose that there are 3 identical and centered vectors with the element (-2,-1,0,1,2); the original calculation (for an even number of views) gives a higher-order covariance of 0, while they should be perfectly correlated. To resolve this issue, for $k = 2m+1$ number of views, each time we take the absolute value of one vector and maintain directionality for the rest of $2m$ vectors to calculate a higher-order correlation, then take the average of all of them. Our empirical findings have shown that this approach is robust in all scenarios.

Eq 1 calculates the higher-order correlation between vectors, but if the higher-order correlation between matrices (e.g. each matrix is an omics dataset) needs to be calculated, we can avoid processing multiple loops to calculate the higher-order correlation for all combinations of features through a different equation. Suppose there are $k$ data in total, denoted by $X_p \in \mathbb{R}^{N\times d_p}, p = 1, 2, .., k$, which are centered with mean 0. for each view, there are $N$ observations and $d_p$ features, then the covariance tensor $C_{1,2,...,k} \in \mathbb{R}^{d_1\times d_2\times ...\times d_k}$ for $k$ views can be denoted by:

$$C_{1,2,...,k} = \begin{cases} |\frac{1}{N}\sum_{j=1}^{N} x_{1j} \circ x_{2j} \circ ... \circ x_{kj}|, \\ \qquad\qquad\qquad\qquad \text{if } k = 2m, m \in \mathbb{N} \\ \frac{1}{k}\sum_{i=1}^{k} C_{1,2,...,|j|,...,k}, \\ \qquad\qquad\qquad\qquad \text{if } k = 2m+1, m \in \mathbb{N}, \end{cases} \tag{2}$$

where

$$C_{1,2,...,|j|,...,k} = \frac{1}{N}|\sum_{i=1}^{N} x_{1i} \circ x_{2i} \circ ... \circ |x_{ji}| \circ ... \circ x_{ki}|, \tag{3}$$

and $\circ$ denotes the outer product. It can be shown that each entry of $C_{1,2,...,k}$, denoted by $C_{1,2,...,k}(j_1, j_2, ..., j_k)$, can also be calculated using Eq. (1), and because of this equivalency, we can avoid processing multiple loops to calculate the higher-order correlation for all combinations of features. The theorem and proof are given below:

**Theorem 1.** *If a covariance tensor $C_{1,2,...,k}$ between $X_1, X_2, ..., X_k$ is calculated by Eq 2, then each entry of $C_{1,2,...,k}$, denoted by $C_{1,2,...,k}(j_1, j_2, ..., j_k)$, is equivalent to:*

$$\rho(x_1(j_1), x_2(j_2), ..., x_k(j_k)) = \begin{cases} \frac{1}{N}|(x_1(j_1) x_2(j_2) ... x_k(j_k))^T \mathbf{1}|, \\ \qquad\qquad\qquad\qquad \text{if } k = 2m, m \in \mathbb{N} \\ \frac{1}{N\cdot k}\sum_{i=1}^{k} |(x_1(j_1) ... |x_i(j_i)| ... x_k(j_k))^T \mathbf{1}|, \\ \qquad\qquad\qquad\qquad \text{if } k = 2m+1, m \in \mathbb{N}, \end{cases} \tag{4}$$

*where $x_j(j_j) \in \mathbb{R}^{N\times 1}$ is the $j_j$th feature of $X_j$.*

*Proof.* If $k = 2m$ is an even number, then

$$\begin{aligned} C_{1,2,...,k} &= \frac{1}{N}|\sum_{i=1}^{N} x_{1i} \circ x_{2i} \circ ... \circ x_{ki}| \\ &= \frac{1}{N}|\Sigma \times_1 X_1^T \times_2 X_2^T \times ... \times_k X_k^T|, \end{aligned} \tag{5}$$

where $\Sigma \in \mathbb{R}^{d_1 \times d_2 \times ... \times d_k}$ is the all-one tensor, and thus each entry of $|C_{1,2,...,k}|$ equals:

$$\begin{aligned} C_{1,2,...,k}(j_1, j_2, ..., j_k) &= \frac{1}{N}|\sum_{i=1}^{N} x_{1i}(j_1)x_{2i}(j_2)...x_{ki}(j_k)| \\ &= \frac{1}{N}|(x_1(j_1)x_2(j_2)...x_k(j_k))^T \mathbf{1}|. \end{aligned} \tag{6}$$

If $k = 2m + 1$ is an odd number, then

$$\begin{aligned} C_{1,2,...,k} &= \frac{1}{k}\sum_{j=1}^{k} |C_{1,2,...,|j|,...k}| \\ &= \frac{1}{N \cdot k}\sum_{j=1}^{k} |\sum_{i=1}^{N} x_{1i} \circ x_{2i} \circ ... \circ |x_{ji}| \circ ... \circ x_{ki}| && (7) \\ &= \frac{1}{N \cdot k}|\Sigma \times_1 X_1^T \times_2 X_2^T \times ... \times_i |X_j^T| \times ... \times X_k^T|, && (8) \end{aligned}$$

and thus each entry of $C_{1,2,...,k}$ equals to:

$$\begin{aligned} C_{1,2,...,k}(j_1, j_2, ..., j_k) &= \frac{1}{N \cdot k}\sum_{j=1}^{k} |\sum_{i=1}^{N} x_{1i}(j_1)x_{2i}(j_2)...x_{ji}(j_i)...x_{ki}(j_k)| \\ &= \frac{1}{N \cdot k}\sum_{j=1}^{k} |(x_1(j_1)...|x_j(j_i)|...x_k(j_k))^T \mathbf{1}|. \end{aligned} \tag{9}$$

□

### S2 Appendix: Proof of Theorem 1

**Theorem 2.** *Let $C_{S_m(j)}$ be the non-negative covariance tensor of view $(m_1, ..., m_m) \in S_m(j)$ such that $C_{S_m(j)} \in \mathbb{R}^{d_{m_1} \times d_{m_2} \times ... \times d_{m_m}}$, If the optimization goal is formulated as follows:*

$$\begin{aligned} \arg\max_{h_1,h_2,...,h_k} \; & a_{k,1}\rho_k(1)^2 + \sum_{j=1}^{\binom{k}{k-1}} a_{k-1,j}\rho_{k-1}(j)^2 + ... \\ & + \sum_{j=1}^{\binom{k}{3}} a_{3,j}\rho_3(j)^2 + \sum_{j=1}^{\binom{k}{2}} a_{2,j}\rho_2(j)^2 \\ \text{s.t.} \quad & h_j^T h_j = 1, j = 1, 2, ..., k, \end{aligned} \tag{10}$$

*where $\rho_m(j) = C_{S_m(j)} \times_1 h_{m_1} \times ... \times_m h_{m_m}$ for all $m = 1, 2, ..., k$, then the optimization problem above is equivalent to the following:*

$$\arg\min_{h_1,h_2,...,h_k} a_{k,1}||C_{S_k(1)} - \hat{C}_{S_k(1)}||_F^2$$
$$+ \sum_{j=1}^{\binom{k}{k-1}} a_{k-1,j}||C_{S_{k-1}(j)} - \hat{C}_{S_{k-1}(j)}||_F^2$$
$$+ ... + \sum_{j=1}^{\binom{k}{3}} a_{3,j}||C_{S_3(j)} - \hat{C}_{S_3(j)}||_F^2$$
$$+ \sum_{j=1}^{\binom{k}{2}} a_{2,j}||C_{S_2(j)} - \hat{C}_{S_2(j)}||_F^2$$
$$\text{s.t. } h_j^T h_j = 1, j = 1, 2, ..., k, \tag{11}$$

*where* $\hat{C}_{S_m(j)} = \rho_m(j) h_{m_1} \circ h_{m_2} \circ ... \circ h_{m_m}$ *is the rank-1 approximation of* $C_{S_m(j)}$.

*Proof.* Based on the formulation above, the augmented Lagrangian of the optimization is given by:

$$f = a_{k,1}||C_{S_k(1)} - \hat{C}_{S_k(1)}||_F^2$$
$$+ \sum_{j=1}^{\binom{k}{k-1}} a_{k-1,j}||C_{S_{k-1}(j)} - \hat{C}_{S_{k-1}(j)}||_F^2$$
$$+ ... + \sum_{j=1}^{\binom{k}{3}} a_{3,j}||C_{S_3(j)} - \hat{C}_{S_3(j)}||_F^2$$
$$+ \sum_{j=1}^{\binom{k}{2}} a_{2,j}||C_{S_2(j)} - \hat{C}_{S_2(j)}||_F^2 + \sum_{j=1}^{k} \alpha_j((||h_j||_2^2 - 1)$$
$$\text{s.t. } h_j^T h_j = 1, j = 1, 2, ..., k \tag{12}$$

Taking the derivative of $f$ with respect to $\rho_m(j), m = 1, 2, ..., k$ and $j = 1, 2, ..., \binom{k}{m}$, and denote $i_{m_s}$ the feature index of the $m_s$-th view ($s = 1, 2, ..., m$), yields the following:

$$\sum_{i_{m_1},...,i_{m_s}} C_{S_m(j)}(i_{m_1}, ..., i_{m_m}) \prod_{m_s \in S_m(j)} h_{m_s}(i_{m_s})$$
$$- \rho_m(j) \sum_{i_{m_1},...,i_{m_s}} \prod_{m_s \in S_m(j)} h_{m_s}(i_{m_s})^2$$
$$= 0 \tag{13}$$

Taking the derivative of $f$ with respect to $\alpha_s, s = 1, 2, ..., k$ yields:

$$\sum_{i_s} h_s(i_s)^2 = 1 \tag{14}$$

then combine the above two equations, we have:

$$\rho_m(j) = \sum_{i_{m_1},...,i_{m_m}} C_{S_m(j)}(i_{m_1}, ..., i_{m_m}) \prod_{m_s \in S_m(j)} h_{m_s}(i_{m_s})$$
$$= C_{S_m(j)} \times_1 h_{m_1} \times_2 h_{m_2} \times ... \times h_{m_m} \tag{15}$$

With the above result, we can finally derive the following:

$$
\begin{aligned}
& a_{k,1}||C_{S_k(1)} - \hat{C}_{S_k(1)}||_F^2 + \sum_{j=1}^{\binom{k}{k-1}} a_{k-1,j}||C_{S_{k-1}(j)} - \hat{C}_{S_{k-1}(j)}||_F^2 \\
& + ... + \sum_{j=1}^{\binom{k}{3}} a_{3,j}||C_{S_3(j)} - \hat{C}_{S_3(j)}||_F^2 + \sum_{j=1}^{\binom{k}{2}} a_{2,j}||C_{S_2(j)} - \hat{C}_{S_2(j)}||_F^2 \\
& = a_{k,1}(||C_{S_k(1)}||_F^2 - 2 < C_{S_k(1)}, \hat{C}_{S_k(1)} > + ||\hat{C}_{S_k(1)}||_F^2) \\
& + \sum_{j=1}^{\binom{k}{k-1}} a_{k-1,j}(||C_{S_{k-1}(j)}||_F^2 - 2 < C_{S_{k-1}(j)}, \hat{C}_{S_{k-1}(j)} > \\
& + ||\hat{C}_{S_{k-1}(j)}||_F^2) + ... \\
& + \sum_{j=1}^{\binom{k}{3}} a_{3,j}(||C_{S_3(j)}||_F^2 - 2 < C_{S_3(j)}, \hat{C}_{S_3(j)} > + ||\hat{C}_{S_3(j)}||_F^2) \\
& + \sum_{j=1}^{\binom{k}{2}} a_{2,j}(||C_{S_2(j)}||_F^2 - 2 < C_{S_2(j)}, \hat{C}_{S_2(j)} > + ||\hat{C}_{S_2(j)}||_F^2),
\end{aligned} \tag{16}
$$

where $< C_{S_m(j)}, \hat{C}_{S_m(j)} >$ is the inner product between two tensors, and $< C_{S_m(j)}, \hat{C}_{S_m(j)} >$ is evaluated as:

$$
\begin{aligned}
& < C_{S_m(j)}, \hat{C}_{S_m(j)} > = \sum_{i_{m_1},...,i_{m_m}} C_{S_m(j)}(i_{m_1}, ..., i_{m_m}) \hat{C}_{S_m(j)}(i_{m_1}, ..., i_{m_m}) \\
& = \sum_{i_{m_1},...,i_{m_m}} C_{S_m(j)}(i_{m_1}, ..., i_{m_m}) \rho_m(j) h_{m_1}(i_{m_1}) \cdot ... \cdot h_{m_m}(i_{m_m}) \\
& = \rho_m(j) \sum_{i_{m_1},...,i_{m_m}} C_{S_m(j)}(i_{m_1}, ..., i_{m_m}) h_{m_1}(i_{m_1}) \cdot ... \cdot h_{m_m}(i_{m_m}) \\
& = \rho_m(j)^2
\end{aligned} \tag{17}
$$

Since $h_j, j = 1, 2, ..., k$ all have the unit norm, combining the equation above with equation 16 we obtain:

$$
\begin{aligned}
& a_{k,1}||C_{S_k(1)} - \hat{C}_{S_k(1)}||_F^2 + \sum_{j=1}^{\binom{k}{k-1}} a_{k-1,j}||C_{S_{k-1}(j)} - \hat{C}_{S_{k-1}(j)}||_F^2 \\
& + ... + \sum_{j=1}^{\binom{k}{3}} a_{3,j}||C_{S_3(j)} - \hat{C}_{S_3(j)}||_F^2 + \sum_{j=1}^{\binom{k}{2}} a_{2,j}||C_{S_2(j)} - \hat{C}_{S_2(j)}||_F^2 \\
& = a_{k,1}(||C_{S_k(1)}||_F^2 - 2\rho_k(1)^2 + \rho_k(1)^2) \\
& + \sum_{j=1}^{\binom{k}{k-1}} a_{k-1,j}(||C_{S_{k-1}(j)}||_F^2 - 2\rho_{k-1}(j)^2 + \rho_{k-1}(j)^2) \\
& + ... + \sum_{j=1}^{\binom{k}{3}} a_{3,j}(||C_{S_3(j)}||_F^2 - 2\rho_3(j)^2 + \rho_3(j)^2) \\
& + \sum_{j=1}^{\binom{k}{2}} a_{2,j}(||C_{S_2(j)}||_F^2 - 2\rho_2(j)^2 + \rho_2(j)^2) \\
& = (a_{k,1}||C_{S_k(1)}||_F^2 + \sum_{j=1}^{\binom{k}{k-1}} a_{k-1,j}||C_{S_{k-1}(j)}||_F^2 + ... \\
& + \sum_{j=1}^{\binom{k}{3}} a_{3,j}||C_{S_3(j)}||_F^2 + \sum_{j=1}^{\binom{k}{2}} a_{2,j}||C_{S_2(j)}||_F^2) \\
& - [\rho_k(1)^2 + \sum_{j=1}^{\binom{k}{k-1}} a_{k-1,j}\rho_{k-1}(j)^2 + ... + \sum_{j=1}^{\binom{k}{3}} a_{3,j}\rho_3(j)^2 \\
& + \sum_{j=1}^{\binom{k}{2}} a_{2,j}\rho_2(j)^2] \\
& = \text{constant} - [\rho_k(1)^2 + \sum_{j=1}^{\binom{k}{k-1}} a_{k-1,j}\rho_{k-1}(j)^2 + ... \\
& + \sum_{j=1}^{\binom{k}{3}} a_{3,j}\rho_3(j)^2 + \sum_{j=1}^{\binom{k}{2}} a_{2,j}\rho_2(j)^2] \qquad (18)
\end{aligned}
$$

Therefore, to minimize equation 11, we need to maximize equation 10, and thus the equivalency holds.

□

### S3 Appendix: Sparse Generalized Tensor Canonical Correlation Analysis (SGTCCA)

**Algorithm 1:** Sparse Generalized Tensor Canonical Correlation Analysis

**Input**:Multi-omics data $X_j \in \mathbb{R}^{N_j \times d_j}$ $(j = 1, 2, ..., k)$, number of sub-samples $s$, proportion of common features $p_c$, proportion of distinct features $p_d$, number of correlation component $l$ ;

**Initialize**: i = 1;

**Iterate the following step until i = s**;

(1) Calculate covariance tensors/matrices of interest $C_1, ..., C_l$;

(2) Calculate covariance density vectors $c_j \in \mathbb{R}^{d_j \times 1}, j = 1, 2, ..., k$ for each molecular profile ;

(3) Rank features based on covariance density, and select the top $d_j \cdot p_c$ features from $j$th view based on density vectors, denoted by $S_{j,c}$;

(4) Randomly select $d_j \cdot p_d$ additional features from $j$th view based on probability vectors $p_j(j = 1, 2, ..., k)$, denoted by $S_{j,d}$, let $S_j = S_{j,c} \cup S_{j,d}$ ;

(5) Implement GTCCA (Algorithm 2 in the main text) in subsampled data $X_j^{(S_j)}$, extract sparse canonical weight matrices $h_j^(i) \in \mathbb{R}^{d_j \times 1}$ (features that are not subsampled received 0 canonical weight) ;

(6) i = i + 1 ;

Concatenate the canonical weights from each sub-sample to the canonical weight matrix $H_j \in \mathbb{R}^{d_j \times s}, j = 1, 2, ..., k$ ;

**Output**: Canonical weight matrices $H_j \in \mathbb{R}^{d_j \times s}, j = 1, 2, ..., k$.

### S4 Appendix: Biased Subsampling Covariance Density

In step (2) of Algorithm 1, the covariance density is defined as follows:

**Definition 1.** *Let $C_{1,2,...,k} \in \mathbb{R}^{d_1 \times d_2 \times ... \times d_k}$ be a tensor with $k$ mode (in covariance tensor, it is equivalent to $k$ views), then the tensor density with respect to mode $j$, denoted by $c_j \in \mathbb{R}^{d_j \times 1}$ is defined as:*

$$c_j = \sum_{i_1,...,i_{j-1},i_{j+1},...,i_k} C_{1,2,...,k}(i_1, ..., i_{j-1}, \cdot, i_{j+1}, ..., i_k). \tag{19}$$

In general, it means that we sum over all other modes except for the mode $j$. The definition above only works for one single tensor when multiple tensors (with shared dimension) are considered, the subsampling density vectors from each tensor need to be aggregated, which is given by the following definition:

**Definition 2.** *Suppose $c_j^{(i)}, j = 1, 2, ..., k, i = 1, 2, ..., l$ ($l$ is the total number of correlation structures) are the density vector for mode $j$ with respect to the $i$th tensor/matrix, then the overall subsampling density vector for the $j$th mode is given by:*

$$c_j = \sum_{i=1}^{l} \frac{(c_j^{(i)})^2}{||c_j^{(i)})^2||_2^2}, \tag{20}$$

where $|| \cdot ||_2$ is defined as the L2 norm of the vector. This formulation ensures that each tensor/matrix contributes equally to the subsampling density vector, and taking the square instead of direct summation further shrinks the density value for features with low covariance density.

As shown in Definition 1, the calculation of the covariance density involves $C_{1,2,...,k}$, which is the full covariance tensor. In practice, this is computationally infeasible. Therefore, we use the unbiased subsampling technique to approximate the covariance density instead, and its definition is as follow:

**Definition 3.** *Let $c_{j,i}^{(sub)} \in \mathbb{R}^{d_j \times 1}, i = 1, 2, ..., s$ be the covariance density vector for the $i$th subsample with only a random subset of features' covariance density value being calculated. To approximate the density vector of the $j$th mode, we take the mean of all subsampled density vectors $c_{j,i}^{(sub)}$ as follows:*

$$c_j = \frac{1}{s}\sum_{i=1}^{s} c_{j,i}^{(sub)}, \tag{21}$$

*where s is the total number of subsampled density vectors, and the mean is calculated by summing all the vectors element-wise and dividing by the number of nonzero entries.*

### S5 Appendix: Network Analysis and Network Pruning Algorithm

**Algorithm 2:** Network Construction and Pruning Algorithm

**Input**: Canonical weight matrices $H_j \in \mathbb{R}^{d_j \times s}, j = 1, 2, ..., k$, Pearson's correlation matrix $\Sigma \in \mathbb{R}^{(\sum_{j=1}^{k} d_j)\times(\sum_{j=1}^{k} d_j)}$, correlation threshold $\rho_{threshold}$ ;

**Initialize**: i = 1 ;

(1) Normalize and concatenate $H_j$ for all $j = 1, 2, ..., k$, and obtain $H \in \mathbb{R}^{(\sum_{j=1}^{k} d_j)\times s}$

(2) **Iterate over the following steps until i = s**

(i) Calculate the adjacency matrix $M = [H(i), H(i)]$, where $[\cdot;\cdot,\cdot]$ is the outer product of the matrix, $H(i)$ is the $i$th column of $H$, and $M \in \mathbb{R}^{(\sum_{j=1}^{k} d_j)\times(\sum_{j=1}^{k} d_j)}$;

(ii) Prune $M$ with PageRank algorithm and NetSHy summarization score and obtain sub-networks $M_{(sub)} \in \mathbb{R}^{(\sum_{j=1}^{k} d_j^{(sub)})\times(\sum_{j=1}^{k} d_j^{(sub)})}$ ;

(iii) Filtering edges from $M_{(sub)}$ with correlation matrix $\Sigma$ and threshold $\rho_{threshold}$ ;

(iv) i = i + 1;

**Output**: Final subnetwork $M_{(sub)}$.

The network pruning algorithm in Step 2 (ii) of the Algorithm 2 is given as follows:

- Calculate the PageRank score for all molecular features in the global network and rank them according to the PageRank score.
- Start from minimally possible network size $m_1$, iterate the following steps until reaching the maximally possible network size $m_2$ (defined by users):
  - Add one more molecular feature into the network based on node ranking, then calculate the NetSHy/PCA summarization score (PC1 - PC3) for this updated network.
  - Calculate the correlation between this network summarization score and phenotype for all possible network sizes $i \in [m_1, m_2]$, and only use the PC with the highest (determined by absolute value) w.r.t. phenotype, define this correlation as $\rho_{(i,pheno)}$, where $i$ stands for the current network size.
- Identify network size $m_*$ ($m_* \in [m_1, m_2]$) with $\rho_{(m_*,pheno)}$ being the maximally possible summarization score correlation w.r.t. phenotype (determined by absolute value).
- Treat $m_*$ as the new baseline network size, let $\rho_{(m_*,i)}$ be the correlation of summarization score between network with size $m_*$ and network with size $i$. Define $x$ to be the network size ($x \in [m_*, m_2]$), such that $x = \max\{i|(i \in [m_*, m_2])\&(|\rho_{(m_*,i)}| > 0.8)\}$.
- Between the size of the network of $m$ and $x$, the optimal size of the network $m_{opt}$ is defined as the maximum size of the network such that $|\rho_{m_{(opt,pheno)}}| \geq 0.9 \cdot |\rho_{(m,pheno)}|$.

### S6 Appendix: Additional Detail of Simulation Study Setup

In the simulation study, suppose $X_1, X_2, X_3$ are 3 omics data and $Y$ is the phenotype data, then the equation for data generation is as follows:

$$
\begin{aligned}
X_1 &= l_1 \cdot b_{1,1}^T + l_2 \cdot b_{2,1}^T + l_3 \cdot b_{3,1}^T + l_5 \cdot b_{5,1}^T \\
&\quad + l_8 \cdot b_{8,1}^T + l_9 \cdot b_{9,1}^T + l_{10} \cdot b_{10,1}^T + \alpha \cdot E_1 \\
X_2 &= l_1 \cdot b_{1,2}^T + l_2 \cdot b_{2,2}^T + l_4 \cdot b_{4,2}^T + l_6 \cdot b_{6,2}^T \\
&\quad + l_8 \cdot b_{8,2}^T + l_9 \cdot b_{9,2}^T + l_{11} \cdot b_{11,2}^T + \alpha \cdot E_2 \\
X_3 &= l_1 \cdot b_{1,2}^T + l_3 \cdot b_{3,3}^T + l_4 \cdot b_{4,3}^T + l_7 \cdot b_{7,3}^T \\
&\quad + l_8 \cdot b_{8,3}^T + l_{10} \cdot b_{10,3}^T + l_{11} \cdot b_{11,3}^T + \alpha \cdot E_3 \\
Y &= \frac{1}{7}\sum_{i=1}^{7} \sigma_i l_i + \beta \cdot E_4
\end{aligned} \tag{22}
$$

Where $\alpha$ and $\beta$ represent the strength of the noise, $\sigma_i = 0, 1$ are the indicators determined by the simulation case (for example, in case 1, $\sigma_i = 1$ for all $i = 1, 2, ..., 7$). In summary, the simulated datasets have 3 types of blocks: (1) signal blocks representing all phenotype-specific correlation structures, which are given by latent factors $l_1, ..., l_7$; (2) non-phenotype-specific blocks representing all non-phenotype-specific correlation structures, which are given by latent factors $l_8, ..., l_{11}$; (3) background noise without any correlation structure, which is given by randomly permuted multi-omics data $E_1, E_2$, $E_3$, and $E_4$. Then the data-generating process is given as follows:

- Simulate 11 latent factors $l_1, l_2, ..., l_{11} \in \mathbb{R}^{N \times 1}$ following multivariate normal distribution $\text{MVN}(0, \Sigma)$, where $\Sigma$ is the identity matrix $11 \times 11$, and each latent factor represents a particular correlation structure.
- Simulate blockwise weight vectors $b_{i,j} \in \mathbb{R}^{1 \times 1000}, i = 1, 2, ..., 11; j = 1, 2, 3$ with nonzero entries corresponding to signal features, and zero entries corresponding to noise features. Nonzero entries from $b_{i,j}$ are calculated as $(2x - 1) \cdot y$, where $x$ (directionality) follows Bernoulli distribution with probability 0.5, and $y$ (magnitude) follows a uniform distribution with the bound of (0.4, 0.6).
- Simulate random noise $E_j \in \mathbb{R}^{N \times d_j}, j = 1, 2, 3, 4$ with each entry following the standard normal distribution.
- Simulate data based on the data generation formula in Eq 22.

We imposed weaker noise on the omics data and stronger noise on the simulated phenotype data to test the noise tolerance of each method. In addition, there are 3 omics data settings we're evaluating (different from the simulation case for correlation structure): (1) Latent factors are simulated with multivariate normal distribution, (2) Latent factors are simulated with highly right-skewed distribution based on Fleishman power transformation, (3) Latent factors are simulated with multivariate normal distribution, but stronger noise is enforced to the omics data.

We compare the performance of our novel pipeline to other multi-omics network analysis methods that produce an adjacency matrix with 20 replications: (1) SGTCCA-Net with only the higher order 4-way correlation structure;(2) Best SmCCNet outcome by various combinations of scaling factors and sparsity levels, (3) Best DIABLO outcome with different levels of sparsity and scaling factors. All these methods have adjacency matrices available for performance evaluation. The performance is evaluated at the node level, and a node is predicted positive if its maximal connection to other nodes in the adjacency matrix passes a certain threshold, which is consistent with SmCCNet evaluation, and the AUC score can be calculated through checking prediction result with a series of the threshold value. By "best" for SmCCNet and DIABLO, we mean running SmCCNet and DIABLO with different combinations of parameters, evaluating the model performance, and only keeping the best-performing AUC score.

#### S1 Table: Simulation Setup Table

| Omics Data Scenarios | Case | Sample Size | Signal Latent Factors | Noise Level |
|---|---|---|---|---|
| Normal Latent Factors: MVN$(0,\Sigma)$ | Case 1 | 100 & 200 | $l_1 - l_7$ | $\alpha = \beta = 0.2$ |
| | Case 2 | 100 & 200 | $l_1$ | $\alpha = \beta = 0.2$ |
| | Case 3 | 100 & 200 | $l_2 - l_7$ | $\alpha = \beta = 0.2$ |
| Highly Right-skewed Latent Factors: Skewness = 2, Kirtosis = 5 | Case 1 | 100 & 200 | $l_1 - l_7$ | $\alpha = \beta = 0.2$ |
| | Case 2 | 100 & 200 | $l_1$ | $\alpha = \beta = 0.2$ |
| | Case 3 | 100 & 200 | $l_2 - l_7$ | $\alpha = \beta = 0.2$ |
| Noisy Omics Data: MVN$(0,\Sigma)$ | Case 1 | 100 & 200 | $l_1 - l_7$ | $\alpha = 0.5, \beta = 0.2$ |
| | Case 2 | 100 & 200 | $l_1$ | $\alpha = 0.5, \beta = 0.2$ |
| | Case 3 | 100 & 200 | $l_2 - l_7$ | $\alpha = 0.5, \beta = 0.2$ |

Table 1: Summary table of all the simulation scenarios. Starting with 3 omics data simulation settings: (1) normal latent factors, (2) highly right-skewed latent factors, and (3) noisy omics data. Each of these settings has 3 cases: case 1 with all phenotype-specific correlation structures; case 2 with only 4-way phenotype-specific correlation structure; and (3) case 3 with 3-way and pairwise phenotype-specific correlation structure. All these cases will be evaluated based on sample sizes of 100 and 200. $\Sigma$ represents a diagonal matrix with diagonal values to be 1.

### S2 Table: Top Network Higher-order Correlation for TCGA Breast Cancer Network

| Type | Gene | miRNA | Protein | Higher-order Correlation |
|---|---|---|---|---|
| 4-way correlation | CLEC4C | hsa-mir-3133 | KDR | 3.938 |
| 4-way correlation | TIMD4 | hsa-mir-3133 | KDR | 3.823 |
| 4-way correlation | SPIB | hsa-mir-3133 | KDR | 3.718 |
| 4-way correlation | LAG3 | hsa-mir-3133 | KDR | 3.479 |
| 4-way correlation | HLA-F | hsa-mir-3133 | KDR | 3.472 |
| 4-way correlation | IL32 | hsa-mir-3133 | KDR | 3.426 |
| 4-way correlation | PATL2 | hsa-mir-3133 | KDR | 3.422 |
| 4-way correlation | AIM2 | hsa-mir-3133 | KDR | 3.361 |
| 4-way correlation | BEND4 | hsa-mir-3133 | KDR | 3.345 |
| 4-way correlation | PSMB9 | hsa-mir-3133 | KDR | 3.268 |
| gene-mirna-phenotype | TMC8 | hsa-mir-142 | - | 1.240 |
| gene-mirna-phenotype | CD48 | hsa-mir-142 | - | 1.196 |
| gene-mirna-phenotype | TMC8 | hsa-mir-150 | - | 1.194 |
| gene-mirna-phenotype | IKZF1 | hsa-mir-142 | - | 1.192 |
| gene-mirna-phenotype | SELL | hsa-mir-142 | - | 1.188 |
| gene-mirna-phenotype | CCR7 | hsa-mir-142 | - | 1.187 |
| gene-mirna-phenotype | IRF8 | hsa-mir-142 | - | 1.185 |
| gene-mirna-phenotype | TMC8 | hsa-mir-146a | - | 1.183 |
| gene-mirna-phenotype | ARHGAP9 | hsa-mir-142 | - | 1.183 |
| gene-mirna-phenotype | CD3E | hsa-mir-150 | - | 1.182 |
| gene-protein-phenotype | TMC8 | - | PCNA | 1.172 |
| gene-protein-phenotype | ARHGAP9 | - | PCNA | 1.141 |
| gene-protein-phenotype | CD48 | - | PCNA | 1.120 |
| gene-protein-phenotype | TBC1D10C | - | PCNA | 1.116 |
| gene-protein-phenotype | LAT | - | PCNA | 1.110 |
| gene-protein-phenotype | RASGRP2 | - | PCNA | 1.107 |
| gene-protein-phenotype | TMIGD2 | - | PCNA | 1.105 |
| gene-protein-phenotype | MAP4K1 | - | PCNA | 1.104 |
| gene-protein-phenotype | IRF8 | - | PCNA | 1.103 |
| gene-protein-phenotype | IKZF1 | - | PCNA | 1.102 |
| mirna-protein-phenotype | - | hsa-mir-3133 | KDR | 1.298 |
| mirna-protein-phenotype | - | hsa-mir-142 | PCNA | 1.045 |
| mirna-protein-phenotype | - | hsa-mir-146a | PCNA | 0.967 |
| mirna-protein-phenotype | - | hsa-mir-150 | PCNA | 0.964 |
| mirna-protein-phenotype | - | hsa-mir-155 | PCNA | 0.925 |
| mirna-protein-phenotype | - | hsa-mir-3692 | PCNA | 0.853 |
| mirna-protein-phenotype | - | hsa-mir-142 | CCNB1 | 0.835 |
| mirna-protein-phenotype | - | hsa-mir-146a | CCNB1 | 0.827 |
| mirna-protein-phenotype | - | hsa-mir-155 | CCNB1 | 0.819 |
| mirna-protein-phenotype | - | hsa-mir-150 | CCNB1 | 0.801 |

Table 2: Top 10 network higher-order correlation of TCGA breast cancer data of different correlation structures.

### S3 Table: Top Network Higher-order Correlation for COPDGene Network

| Type | Gene | Protein | Metabolite | Higher-order Correlation |
|---|---|---|---|---|
| 4-way correlation | MTCO1P12 | Tyrosine-protein kinase Yes | C-glycosyltryptophan | 1.216 |
| 4-way correlation | MTCO1P12 | Tyrosine-protein kinase Yes | N-acetylneuraminate | 1.151 |
| 4-way correlation | HMGB1P5 | Tyrosine-protein kinase Yes | N-acetylneuraminate | 0.934 |
| 4-way correlation | HMGB1P5 | Tyrosine-protein kinase Yes | C-glycosyltryptophan | 0.922 |
| 4-way correlation | MTCO1P12 | Tyrosine-protein kinase Yes | glycodeoxycholate | 0.913 |
| 4-way correlation | RPL7P9 | Tyrosine-protein kinase Yes | C-glycosyltryptophan | 0.878 |
| 4-way correlation | RPL7P9 | Tyrosine-protein kinase Yes | N-acetylneuraminate | 0.872 |
| 4-way correlation | MTCO1P12 | Tyrosine-protein kinase Yes | 5-acetylamino-6-amino-3-methyluracil | 0.867 |
| 4-way correlation | MTCO1P12 | Tyrosine-protein kinase Yes | N-acetyl-1-methylhistidine* | 0.818 |
| 4-way correlation | HK3 | Plasma serine protease inhibitor | 1-stearoyl-2-oleoyl-GPI (18:0/18:1)* | 0.762 |
| gene-protein-phenotype | S100A9 | Matrix metalloproteinase-9 | - | 0.302 |
| gene-protein-phenotype | GCA | Matrix metalloproteinase-9 | - | 0.301 |
| gene-protein-phenotype | GLT1D1 | Matrix metalloproteinase-9 | - | 0.297 |
| gene-protein-phenotype | MGAM2 | Matrix metalloproteinase-9 | - | 0.296 |
| gene-protein-phenotype | BCL6 | Matrix metalloproteinase-9 | - | 0.296 |
| gene-protein-phenotype | CSF3R | Matrix metalloproteinase-9 | - | 0.296 |
| gene-protein-phenotype | SLC11A1 | Matrix metalloproteinase-9 | - | 0.293 |
| gene-protein-phenotype | LIN7A | Matrix metalloproteinase-9 | - | 0.290 |
| gene-protein-phenotype | ALOX5 | Matrix metalloproteinase-9 | - | 0.290 |
| gene-protein-phenotype | SULT1B1 | Matrix metalloproteinase-9 | - | 0.290 |
| gene-metabolite-phenotype | TTTY15 | - | 1-stearoyl-2-linoleoyl-GPI (18:0/18:2) | 0.226 |
| gene-metabolite-phenotype | MGAM2 | - | 1-stearoyl-2-oleoyl-GPI (18:0/18:1)* | 0.211 |
| gene-metabolite-phenotype | HK3 | - | 1-stearoyl-2-oleoyl-GPI (18:0/18:1)* | 0.208 |
| gene-metabolite-phenotype | MGAM2 | - | 1-(1-enyl-stearoyl)-GPE (P-18:0)* | 0.204 |
| gene-metabolite-phenotype | CMTM2 | - | X - 12544 | 0.196 |
| gene-metabolite-phenotype | TTTY15 | - | 1-stearoyl-2-docosahexaenoyl-GPE (18:0/22:6)* | 0.196 |
| gene-metabolite-phenotype | PRKY | - | 1-stearoyl-2-linoleoyl-GPI (18:0/18:2) | 0.193 |
| gene-metabolite-phenotype | CR1 | - | 1-stearoyl-2-oleoyl-GPI (18:0/18:1)* | 0.191 |
| gene-metabolite-phenotype | S100A9 | - | 1-stearoyl-2-oleoyl-GPI (18:0/18:1)* | 0.189 |
| gene-metabolite-phenotype | NCF4 | - | X - 12544 | 0.185 |
| protein-metabolite-phenotype | - | Tyrosine-protein kinase Yes | C-glycosyltryptophan | 0.289 |
| protein-metabolite-phenotype | - | Neutrophil gelatinase-associated lipocalin | C-glycosyltryptophan | 0.287 |
| protein-metabolite-phenotype | - | Tyrosine-protein kinase Yes | N-acetylneuraminate | 0.283 |
| protein-metabolite-phenotype | - | Fibrinogen gamma chain | iminodiacetate (IDA) | 0.268 |
| protein-metabolite-phenotype | - | D-dimer | iminodiacetate (IDA) | 0.254 |
| protein-metabolite-phenotype | - | Neutrophil gelatinase-associated lipocalin | N-acetylneuraminate | 0.251 |
| protein-metabolite-phenotype | - | MMP-8 | 6-hydroxyindole sulfate | 0.238 |
| protein-metabolite-phenotype | - | MMP-8 | iminodiacetate (IDA) | 0.237 |
| protein-metabolite-phenotype | - | Neutrophil gelatinase-associated lipocalin | 5-acetylamino-6-amino-3-methyluracil | 0.235 |
| protein-metabolite-phenotype | - | Peptide YY | 5-hydroxyhexanoate | 0.233 |

Table 3: Top 10 network higher-order correlation of COPDGene data of different correlation structures.

### S4 Table: Top Metabolite Enrichment Pathways

| Pathway Names | Source | Enriched | Background (All) | P-value |
|---|---|---|---|---|
| Phospholipid Biosynthesis | SMPDB | 4 | 10 (27) | 0.001 |
| Glycerophospholipid metabolism | KEGG | 3 | 8 (55) | 0.007 |
| Glycerophospholipid biosynthesis | Reactome | 3 | 12 (94) | 0.024 |
| Phospholipid metabolism | Reactome | 3 | 12 (104) | 0.024 |
| 27-Hydroxylase Deficiency | SMPDB | 3 | 14 (64) | 0.037 |
| Congenital Bile Acid Synthesis Defect Type III | SMPDB | 3 | 14 (64) | 0.037 |
| Bile Acid Biosynthesis | SMPDB | 3 | 14 (64) | 0.037 |
| Congenital Bile Acid Synthesis Defect Type II | SMPDB | 3 | 14 (64) | 0.037 |
| Cerebrotendinous Xanthomatosis (CTX) | SMPDB | 3 | 14 (64) | 0.037 |
| Zellweger Syndrome | SMPDB | 3 | 14 (64) | 0.037 |

Table 4: Top 10 pathways of metabolite enrichment analysis result from IMPaLa for SGTCCA-Net subnetwork.

### S5 Table: Overlap Molecular Features between FEV1 and FEV1 Percent Predicted Network

| Name | Type | Correlation (FEV1) | P-value (FEV1) | Correlation (FEV1 PP) | P-value (FEV1 PP) |
|---|---|---|---|---|---|
| C-reactive protein | Protein | -0.190 | <0.001 | -0.257 | <0.001 |
| ENSG00000111052 | Gene | -0.167 | <0.001 | -0.220 | <0.001 |
| Complement C5 | Protein | -0.152 | 0.001 | -0.200 | <0.001 |
| 5-hydroxyhexanoate | Metabolite | -0.170 | <0.001 | -0.190 | <0.001 |
| mannose | Metabolite | -0.115 | 0.013 | -0.183 | <0.001 |
| ENSG00000101916 | Gene | -0.149 | 0.001 | -0.175 | <0.001 |
| ENSG00000160883 | Gene | -0.177 | <0.001 | -0.174 | <0.001 |
| Complement C5b-C6 complex | Protein | -0.138 | 0.003 | -0.173 | <0.001 |
| ENSG00000115271 | Gene | -0.133 | 0.004 | -0.169 | <0.001 |
| ENSG00000163220 | Gene | -0.138 | 0.003 | -0.166 | <0.001 |
| ENSG00000173597 | Gene | -0.143 | 0.002 | -0.156 | <0.001 |
| Matrix metalloproteinase-9 | Protein | -0.149 | 0.001 | -0.154 | <0.001 |
| 1-linoleoyl-GPC (18:2) | Metabolite | 0.126 | 0.007 | 0.160 | <0.001 |
| 1-(1-enyl-stearoyl)-GPE (P-18:0)* | Metabolite | 0.151 | 0.001 | 0.174 | <0.001 |
| Plasma serine protease inhibitor | Protein | 0.180 | <0.001 | 0.219 | <0.001 |
| Kallistatin | Protein | 0.158 | <0.001 | 0.223 | <0.001 |

Table 5: Molecular features overlap between FEV1 and FEV1 percent predicted network generated from SGTCCA-Net (where absolute correlation with phenotype >0.15).
